# The extracellular contractile injection system is enriched in environmental microbes and associates with numerous toxins

**DOI:** 10.1101/2020.09.22.308684

**Authors:** Alexander Martin Geller, Inbal Pollin, David Zlotkin, Aleks Danov, Nimrod Nachmias, William B Andreopoulos, Keren Shemesh, Asaf Levy

## Abstract

Bacteria employ toxin delivery systems to exclude bacterial competitors and to infect host cells. Characterization of these systems and the toxins they secrete is important for understanding microbial interactions and virulence in different ecosystems. The extracellular Contractile Injection System (eCIS) is a toxin delivery particle that evolved from a bacteriophage tail. Four known eCIS systems have been shown to mediate interactions between bacteria and their invertebrate hosts, but the broad ecological function of these systems remains unknown. Here, we identify eCIS loci in 1,249 prokaryotic genomes and reveal a striking enrichment of these loci in environmental microbes and absence from mammalian pathogens. We uncovered 13 toxin genes that associate with eCIS from diverse microbes and show that they can inhibit growth of bacteria, yeast or both. We also found immunity genes that protect bacteria from self-intoxication, supporting an antibacterial role for eCIS. Furthermore, we identified multiple new eCIS core genes including a conserved eCIS transcriptional regulator. Finally, we present our data through eCIStem; an extensive eCIS repository. Our findings define eCIS as a widespread environmental prokaryotic toxin delivery system that likely mediates antagonistic interactions with eukaryotes and prokaryotes. Future understanding of eCIS functions can be leveraged for the development of new biological control systems, antimicrobials, and cell-free protein delivery tools.

## Introduction

The extracellular contractile injection system (eCIS; previously termed “PLTS”, phage-like protein-translocation structures) is a cell-free protein delivery system that is prevalent in bacteria and archaea, but its biological function is poorly understood. The eCIS particle resembles the contractile tail of a T4 bacteriophage and is mostly encoded by an operon of 15-28 genes ^1^. This protein complex is 110-120 nm long and includes a baseplate, a sheathed hollow tube that has a needle-like tip (spike) on one side and a cap on the other side, and tail fibers that likely serve to adhere to target cells (Figure S1) ^2,3^. eCIS contraction propels the tube out of the sheath, likely enabling the sharp tip to perforate the target cell membrane. Effectors, which are proteins that are injected by the particle eCIS into target cells upon contraction. These effectors are usually encoded at the 3’ end of the operon ^4–6^. The effectors that have been studied were shown to perform enzymatic activities in the target eukaryotic cell, most of which lead to cell toxicity. eCIS shares structural similarity with other contractile nano weapons such as type VI secretion system (T6SS) and R-type pyocins but differs from these by being extracellular and by injecting effectors into the target cell instead of just perforating it, respectively. Despite the prevalence of eCISs across the microbial world, only four eCIS loci, Afp, AfpX, PVC, and MACs, have been experimentally studied.

The Antifeeding Prophage (Afp) is encoded by *Serratia entomophila* and it is sufficient to cause feeding cessation and death to the New Zealand grass grub pest ^7–10^. A homologous Afp structure, termed AfpX, was described in *Serratia proteamaculans* and demonstrated insecticidal activity against the larvae of two insects ^6^. Effectors of Afp and AfpX have not been confirmed. Photorhabdus Virulence Cassettes (PVC) was characterized in *Photorhabdus* spp. and has an injectable insecticidal activity against wax moth larvae ^5^. Four effectors are associated with PVC and demonstrate cytotoxicity against eukaryotic cells: Plu1690 ^5^, Pnf ^4^, RRSP_Pa_ ^11^, and SepC-like ^5^. The high-resolution structures of Afp and PVC particles were recently solved using cryo-EM and provided excellent maps of the protein interactions within these particles ^2,3^. Unlike the insecticidal activities of Afp, AfpX, and PVC, the metamorphosis-associated contractile structures (MACs) encoded by *Pseudoalteromonas luteoviolacea* plays a mutualistic role. MACs loci produce arrays of ~100 eCIS structures that trigger metamorphosis of the marine tubeworm *Hydroides elegans* hosting the bacteria ^12^. MACs particles inject a nuclease effector that kills eukaryotic cell lines but this activity is not essential for metamorphosis ^13^. Mif1 protein was shown as a cargo inside the MACs tube lumen but was not shown to act as a toxin ^14^. To summarize, eCIS particles interact with different invertebrates and inject effectors that are toxic to eukaryotic cells. To the best of our knowledge, only seven eCIS effectors were experimentally validated as toxic to recipient cells or are known to be injected by eCIS. All known effectors are encoded in gammaproteobacteria.

Here we provide a systematic characterization of eCIS in the microbial world. We identified 1,425 eCIS loci encoded within 1,249 prokaryotic genomes, corresponding to 1.9% of the analyzed microbial genomes. Strikingly, eCIS loci are strongly enriched in environmental microbial taxa from different ecosystems and in microbiomes of specific hosts (plants, specific animals, protists), and are depleted from mammalian pathogens that have been extensively cultured and sequenced. We analyzed the proteins and protein domains of all eCIS loci and identified new core protein domains, including a putative metallopeptidase that associates with toxins and a putative master transcriptional regulator. We further bioinformatically analyzed the fibre proteins of eCIS that confer cell specificity and identified the first group of eCIS particles likely targeting bacteria. We identified a large set of putative toxins (effectors) that are genetically associated with eCIS operons, the majority of which are specific to certain eCIS loci. Through heterologous expression experiments we showed that 13 toxins were capable of killing *E. coli* and/or *S. cerevisiae* cells. In some cases, we were able to experimentally identify immunity genes against that rescued bacteria from self-intoxication by their cognate toxins, supporting the toxin intended activity against bacteria. Finally, we developed eCIStem; an online resource to present our findings that will serve as a valuable resource for the research community. We thus provide here an in-depth ecological and functional characterization of a poorly studied toxin delivery system and its associated toxins.

## Results

### eCIS are encoded by 1.9% and 1.2% of sequenced bacteria and archaea, respectively, with a highly biased taxonomic distribution

First, we were interested in identifying all eCIS loci in a large genomic dataset. We compiled a set of 64,756 microbial isolate genomes retrieved from Integrated Microbial Genomes ^15^. To identify core component homologs from known systems, we searched for genes with known eCIS-associated pfam annotations. To supplement this, we also annotated homologous genes ourselves by searching using Hidden Markov Model (HMM) profiles from a recent publication^1,16^. We defined putative eCIS operons as gene cassettes that included these multiple eCIS core genes in close proximity, and were not bacteriophage, T6SS, or R-type pyocins (Materials and Methods). Overall, we identified eCIS operons encoded in 1,230 (1.9%) bacteria and 19 (1.2%) archaea from our genomic repository (Table S1-S2). We identified two core genes, *Afp8* and *Afp11,* that are present together across 98.7% all eCIS loci and used their protein sequences to construct an eCIS phylogenetic gene tree (Figure 1a). eCIS is scattered across the prokaryotic diversity with presence in 14 bacterial phyla and in one archaeal phylum. The incongruence between this tree and the genomic phylogeny suggests that eCIS undergo HGT frequently, as was proposed before ^1,16^. The previously experimentally characterized eCIS are located within a narrow clade on the eCIS tree, pointing to the possibility that other eCIS particles may play more diverse ecological roles (Figure 1a).

**Figure 1.**
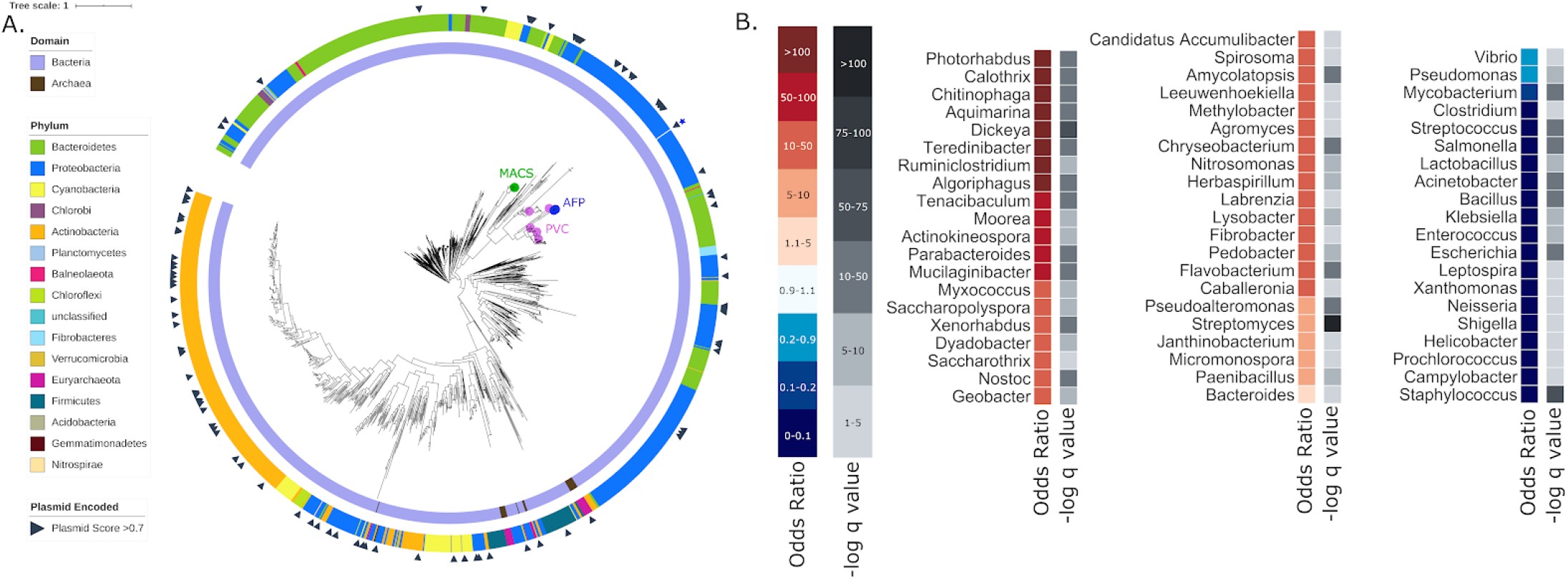
Taxonomic Distribution of eClS-encoding microbes. **A. A phylogenetic tree of eCIS across the microbial world.** eCIS core genes Afp8 and Afp11 from each operon were concatenated, aligned, and used to construct the phylogenetic tree. The Domain and Phylum corresponding to each leaf are indicated in the inner and outer rings, respectively. Scaffolds encoding eCIS were predicted to be plasmids using Deeplasmid and marked with a triangle. Previously experimentally investigated eCIS are marked on their respective leaves (2 o’clock). **B. eCIS distribution in different genera.** We calculated the eCIS distribution across genera using a Fisher exact test. The Odds Ratio represents the enrichment or depletion magnitude, with redder colors representing enrichment, and bluer colors representing depletion. Calculated p values were corrected for multiple testing using FDR to yield minus log10 q values, shown in shades of grey.

Next, we looked for genetic mechanisms that may mediate the eCIS HGT. Using Deeplasmid, a new plasmid prediction tool that we developed, we identified that 7.6% of eCIS are likely plasmid-borne (Figures 1a and S2, Table S3, Materials and Methods). In other cases, we found a clear signature of eCIS operon integration into a specific bacterial chromosome (Figure S3). For example, we identified a likely homologous recombination event between identical tRNA genes, a classical integration site ^17^ (Figure S3b). These genomic integration events and the plasmid-borne eCIS operons shed light on the mechanisms through which eCIS loci have been horizontally propagated in microbial genomes.

### eCIS displays a highly biased taxonomic distribution

Given the propensity of eCIS to transfer between microbes as phylogenetically distant as bacteria and archaea, we were surprised by its scarcity in microbial genomes. We tested if eCIS loci are homogeneously distributed across microbial taxa and found that eCIS are mostly constrained to particular taxa (Figure 1b, Table S4). Strikingly, we found that it is present in 100% (18/18) of *Photorhabdus* genomes in our dataset with 2-5 eCIS operon copies per genome (Fisher exact test, odds ratio = infinity, q value = 2.97e^−28^), 89% of sequenced *Chitinophaga* (odds ratio = 276, q value = 1.69e^−35^), 86% of sequenced *Dickeya* (odds ratio = 211, q value = 3.78e^−52^), and 69% of sequenced *Algoriphagus* (odds ratio = 73, q value = 1.99e^−24^). These genera are known as environmental microbes; *Photorhabdus* is a commensal of entomopathogenic nematodes ^18^, *Chitinophaga* is a soil microbe and a fungal endosymbiont ^19^, *Dickeya* is a plant and pea aphid pathogen ^20,21^, and *Algoriphagus* is an aquatic or terrestrial microbe ^22–26^. In contrast, eCIS is strongly depleted from the most cultured and sequenced genera of Gram-positive and negative human pathogens, including *Staphylococcus, Escherichia, Salmonella, Streptococcus, Acinetobacter,* and *Klebsiella.* Strikingly, within these genera, for which our repository had 18,355 genomes, eCIS was totally absent (odds ratio = 0, q value ≤ 3.86E-16 for each one of these genera), suggesting a very potent purifying selection acting against eCIS integration into these microbial genomes, despite the eCIS operon’s tendency for extensive lateral transfer and its presence in other host-associated systems.

### eCIS presence is highly correlated with specific ecosystems, microbial lifestyles, and microbial hosts

Given the strong eCIS taxonomic bias we identified, we were curious to know if we could further associate eCIS with specific ecological features. To this end, we retrieved metadata available for all sequenced genomes in our repository (Materials and Methods). These traits include the microbial isolation site, ecosystem and habitat, microbial lifestyle and physiology, and the organisms hosting the microbes (Table S5). We calculated the correlation of eCIS presence with certain microbial traits to identify significant enrichments and depletion patterns. This was done using a naïve enrichment test (Fisher exact test) together with a phylogeny-aware test, Scoary ^27^, which is used to correct for the phylogenetic bias of the isolate genomes. Using this test we quantify to what extent the eCIS presence in a genome correlates with a certain trait, independently of the microbial phylogeny (Figure 2). Notably, eCIS is positively correlated with terrestrial and aquatic environments, such as soil, sediments, lakes, and coasts, but is depleted from built environments and food production venues. In terms of microbial lifestyle and physiology, eCISs are enriched in environmental microbes, mostly symbiotic, and are depleted from pathogens (the vast majority of which were isolated from humans). eCISs are enriched in aerobic microbes that dwell in mild and cold temperatures. In general, the eCIS-encoding microbes tend to associate with terrestrial hosts including insects, nematodes, annelids, protists, fungi, and plants, and in aquatic hosts such as fish, sponges, and molluscs. Intriguingly, we detected a strong depletion from bacteria that were isolated from birds and mammals, including humans. Specifically, the operon is depleted from all tissues in which the human microbiome is abundant: oral and digestive systems, skin, and the urogenital tract. However, we detected a mild eCIS enrichment in the human gut commensal *Bacteroides* (Figure 1B) and *Parabacteroidetes* genera. *Bacteroides* was recently reported by the Shikuma group as being eCIS-rich ^28^. Given these findings, we conclude that eCIS is a strongly enriched environmental microbial secretion system. Moreover, to the best of our knowledge, this is the first secretion system that exhibits such a strong genomic enrichment in environmental microbes and a depletion from mammalian and avian microbiomes. Therefore, we hypothesize that this environmental association points to the eCIS biological function and target organism specificity.

**Figure 2.**
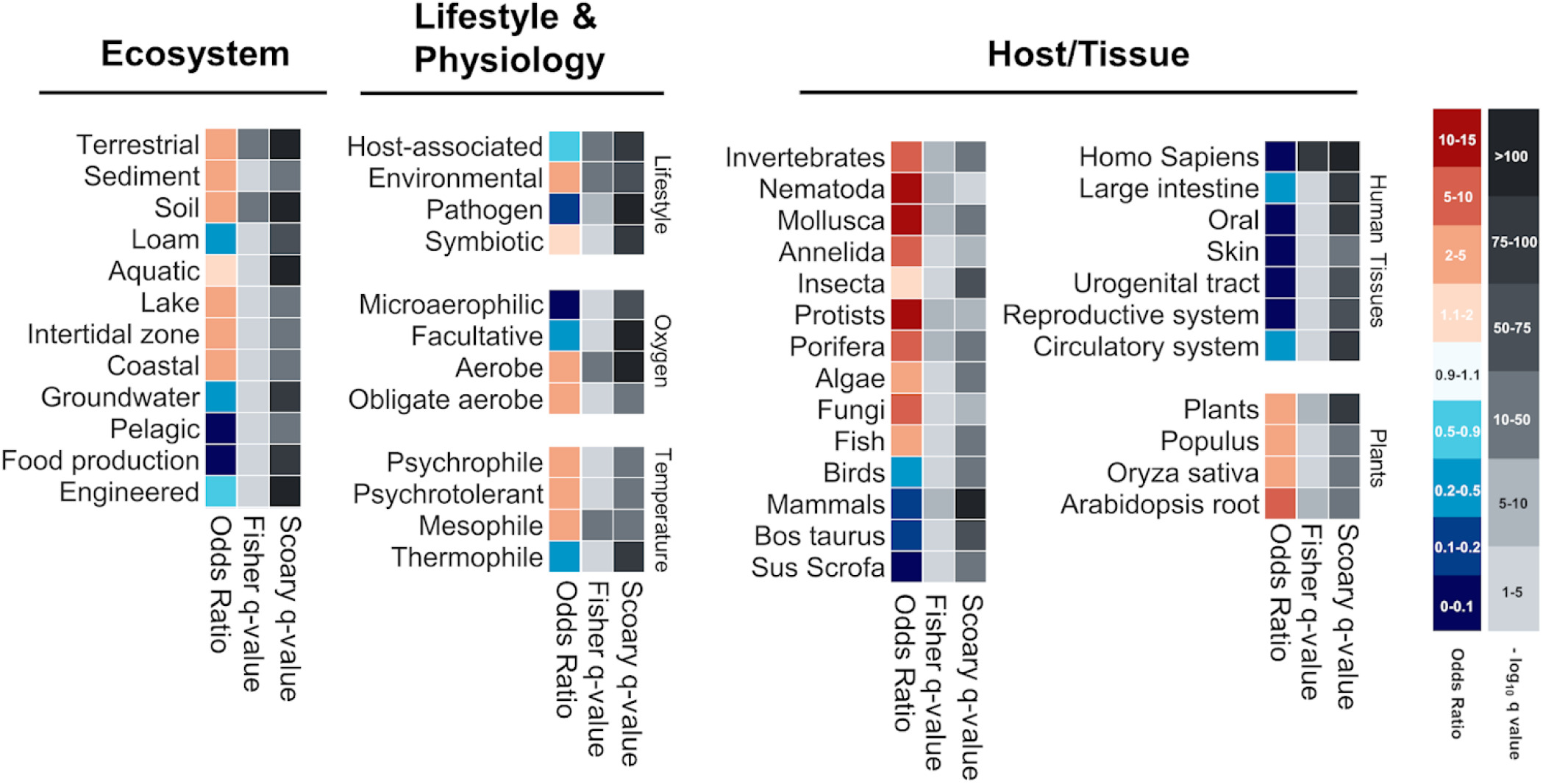
eCIS-encoding microbes’ lifestyle and isolation. A Fisher exact test combined with a modified version of Scoary was used to perform a population-aware analysis of eCIS-encoding microbes’ metadata. The Odds Ratio represents the enrichment or depletion magnitude, with redder colors representing enrichment, and bluer colors representing depletion. The negative log10 of the q-values, shown in shades of grey, are corrected for multiple hypothesis testing. One q-value corresponds to the statistical significance of the Fisher exact test, and the other represents the same for the Scoary pairwise comparison test.

### eCIS tail fibres provide information about target specificity and suggest targeting of bacteria

All previously studied eCIS target invertebrate cells. One of the proteins that likely mediates specific eCIS binding to target cells prior to effector injection is the tail fibre protein that is bound to the baseplate ^29^. This protein is encoded by the *Afp13* gene and has homologues across the different eCIS operons. The Afp13 from *Serratia* spp. was shown to share amino acid sequence similarity with the tail fibre protein of adenovirus, an eukaryotic-targeting virus ^6,10^. We sought to classify the Afp13 homologues in our dataset according to the fiber protein they are most similar to, whether from eukaryote-targeting viruses or from bacteria-targeting viruses (phages) to assign potential eCIS target cells. To do so we compiled a database of all proteins from 192,410 eukaryote-targeting viruses and 12,508 tailed phages and used protein BLAST with 629 Afp13 protein sequences as the query (Table S6, Materials and Methods). We identified Afp13 that have strong matches to eukaryote-targeting viruses, including fibre proteins from Equine Adenovirus 1, Bat mastadenovirus B, and Rhesus adenovirus 60. Interestingly, hits were also detected against the algae-infecting Organic Lake Phycodnavirus, as well as against amoeba-infecting Yasminevirus (Figure S4a,b), suggesting new eCIS eukaryotic targets beyond invertebrates. Our data showed that 99 Afp13 sequences matched eukaryotic-targeting viruses better than they matched phages. However, 16 Afp13 sequences aligned best with phage fibres (Figure S4c-f, Table S6). We therefore propose that in these cases, the eCIS particles likely target bacteria. This notion is supported by the fact that highly-related contractile tail particles, R-type pyocins, have analogous tail fibers that indeed specify host range by binding to specific strains of bacteria ^30,31^.

Notably, only 18% of the Afp13 protein sequences tested share a clear sequence similarity to a known tail fibre from phages or from eukaryotic-targeting viruses (but not to both groups). Therefore, in the majority of the cases, it is challenging to predict the kingdom to which the eCIS target organism belongs solely based on tail fibre protein sequence.

### Characterization of the eCIS known and new core and shell protein domains

Hundreds of eCIS operons were previously defined ^1,16^ but a systematic analysis of the genes that compose the eCIS operons has not been carried out yet. To comprehensively characterize the eCIS repertoire of genes, we compiled the list of Protein Family (pfam) domains across the 1,425 eCIS operons in our database (Materials and Methods). We then performed an enrichment test to identify protein domains that are statistically enriched in eCIS operons in comparison to all other genes in our 64,756 genome dataset (Figure 3a, Table S7). As expected, most of the known core eCIS protein domains are highly enriched and are present in nearly all eCIS loci (Figure 3a and b). These include the domains encoding eCIS tail tube, spike complex, sheath, baseplate, tail terminator protein, and ATP supply (Figure 3a). To identify yet unknown core domains we searched for protein domains that are present in eCIS from at least 10 microbial phyla. By doing so we identified a novel highly conserved eCIS core domain of unknown function 4157 (DUF4157). DUF4157 is found in 13 phyla and has high eCIS specificity (odd ratio = 2614, q value = 0). It contains the Zn-binding motif HExxH which characterizes many metalloproteases ^32–35^. This domain, previously termed PVC-metallopeptidase, was identified as a marker of eCIS based on limited genome analyses ^36,37^ but its role has never been studied in a broad and functional context. By analyzing the eCIS operons and proteins we identified that DUF4157 is strongly associated with a high number of putative eCIS toxins, either as a gene that is adjacent to toxins or in the form of a multidomain protein, in which DUF4157 is found at the N-terminus and the toxin domain is found at the C-terminus (Figure S6). Interestingly, in many cases there is a very long linker sequence between DUF4157 and the various toxin domains, occasionally more than 1000 amino acids long (Figure S6). The role of the DUF4157 domain in eCIS remains to be studied. We hypothesize that it is related to toxin loading, release, maturation, or trafficking inside the target cell.

**Figure 3.**
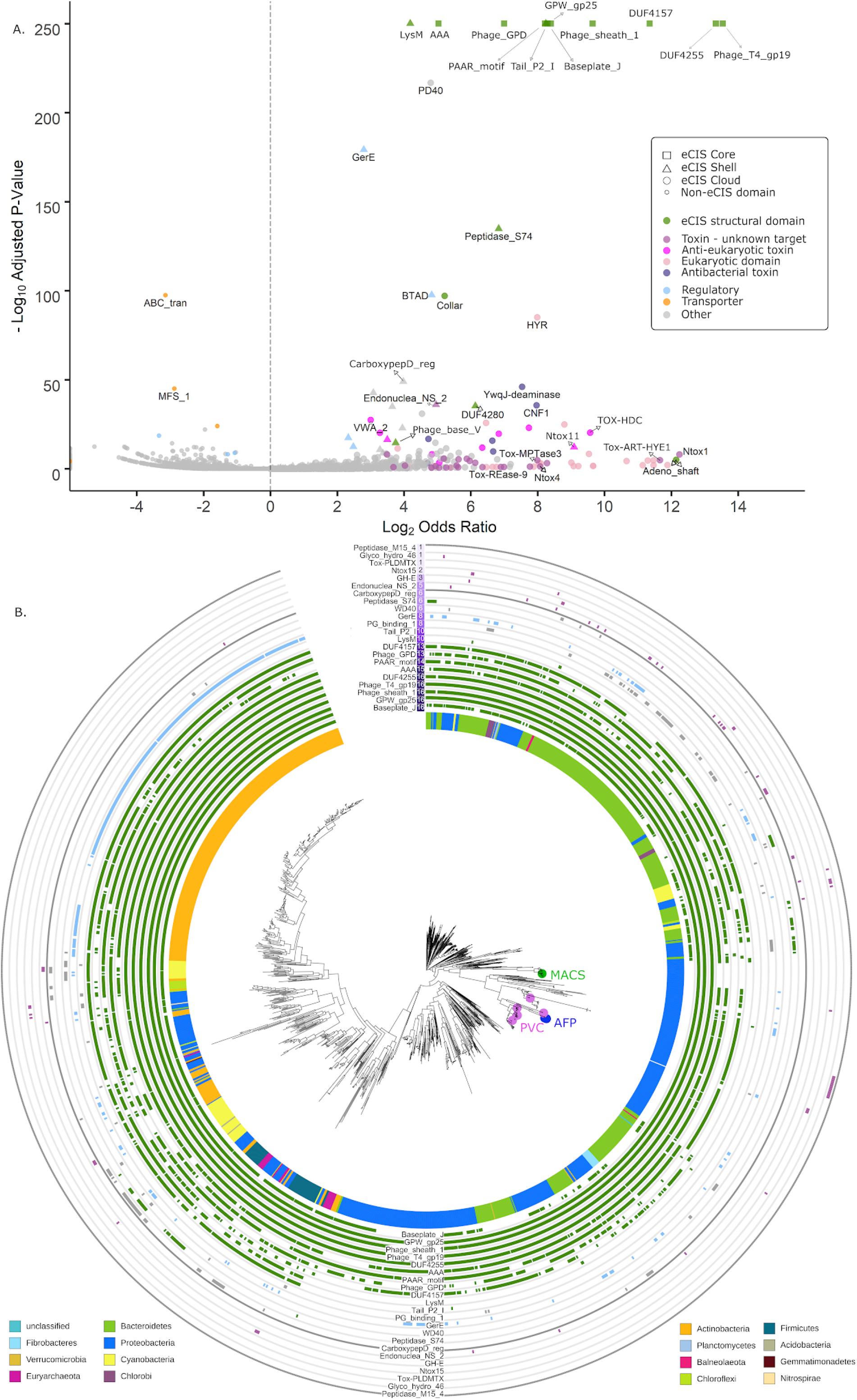
Protein Family (Pfam) domains that are enriched in eCIS. **A. Volcano plot of pfam domain enrichment in eCIS operons.** Fisher Exact test corrected with Benjamini Hochberg procedure. X axis is Log2 of odds ratio, Y axis is negative log10 of corrected P-value. The shape of each point is represented by the number of Phyla containing the Pfam in eCIS operon; >13 is defined “Core”, 4-10 is “Shell”, <4 is “Cloud”. The color of each point represents a functional context of the Pfam domain. The known eCIS core domains encode the tail tube (Phage_T4_gp19), spike complex (PAAR_motif, Phage_GPD), sheath (Phage_sheath_1), baseplate (GPW_gp25, Baseplate_J), tail terminator protein (DUF4255), and ATP supply (AAA). DUF4157 is a novel eCIS core domain., **B. Phylogenetic distribution of eCIS-associated pfams.** At the 12 o’clock and 6 o’clock positions of the tree are labels for select eCIS-associated pfams. Next to each pfam domain name at 12 o’clock is a heatmap quantifying how many phyla a given pfam is found in, along with a ring corresponding to each label (surrounding the circumference of the tree) with colors corresponding to A.

Next, we defined ‘shell’ domains as those that are present in eCIS from 4-10 microbial phyla and likely produce specific eCIS subtypes or are responsible for regulation (Figure 3, Table S7). These shell domains include, for example, LysM (PF01476), Tail_P2_I (PF09684), and the bacterial transcriptional activator domain (BTAD, PF03704). An example of a novel eCIS shell domain is the GerE transcriptional regulator (PF00196) which is a LuxR-type DNA-binding domain. We find GerE located next to 25.8% of the eCIS loci, thereby defining it as the first conserved eCIS transcriptional regulator (Figure 3a and b). In 9% of the eCIS operons containing the GerE transcriptional regulator, a histidine kinase gene was found upstream to the GerE genes (Figure S8). These two genes might constitute a two-component system regulating the eCIS operon. Another novel shell domain is the Peptidase S74 (PF13884). The Peptidase S74 is a known domain of the intramolecular chaperone of the bacteriophage T5 tail fibre ^38^ (Figure S9). eCIS operons containing Peptidase S74 genes did not have annotated tail fibres (*Afp13*), suggesting they might have similar functions.

### A large collection of toxin domains is encoded within eCIS operon genes

We identified numerous toxin domains located next to eCIS core components. Intriguingly, unlike the core and shell domains, most putative toxins are specific to certain eCIS loci (Figure 3b), suggesting rapid evolution and diversification of these toxins as part of the arms race against different target cells. Some of these domains are likely antibacterial toxins, such as: a DNAse with an Ntox15 domain (PF15604) ^39^ and pesticin (PF16754) ^40^, which are encoded by the eCIS of *Lysobacter antibioticus* and *Pseudoalteromonas luteoviolacea,* respectively, supporting the idea that some eCIS are released to target bacteria. Other toxins, as expected, seem to target eukaryotes, e.g.: cycle inhibiting factor (PF16374) that arrest cell cycle ^41^, and Von Willebrand factor type A domain, which is a glycoprotein found in blood plasma (PF00092). In addition, we predict that other eCIS domains serve as eCIS anti-eukaryotic toxin effectors as they are mostly found in eukaryotic proteins (Figure S7), such as hemopexin that is found in plasma proteins (PF00045) ^42^, Annexin (PF00191), and the animal-specific prominin domain (PF05478) ^43^. Eukaryotic domains are common in effectors translocated into eukaryotic cells by various secretion systems ^44–47^. Overall, we identified at least 71 protein domains that are likely toxins and are found next to eCIS core genes.

### Experimental validation of 12 novel eCIS-associated toxins that target bacteria

Next, we decided to test our computational toxin gene predictions. We tested killing of *E. coli* as we predict that certain eCIS particles inject toxins into bacteria. To identify new toxins we first clustered all eCIS proteins based on sequence similarity and annotated core genes (Materials and Methods, Table S8). We selected 20 candidate eCIS toxins based on several features, such as small size, encoding domains (e.g. proteins with DUF4157), and location at the 3’ end of their respective operon ^4–6^. We synthesized and cloned genes of interest under an inducible T7 promoter in pET vectors and transformed these into *E. coli.* We used a known antibacterial toxin, Tse2 ^48^, as a positive control, and two negative controls: a monomeric enhanced GFP gene ^49^ and an eCIS phage tail protein that is not expected to be toxic (Ga0070536_101452). Following serial dilutions of the colonies we could quantify the expressed protein toxicity levels. Out of the 20 genes tested, 12 were found to be toxic to *E. coli* (Figure 4a). We designate these as eCIS-associated toxins (“EATs”) 1-12. Based on sensitive sequence similarity search we identified that the EATs likely encode various molecular functions, such as ADP ribosylation, peptidase, NAD+ phosphorylase, endonuclease, peptidoglycan hydrolysis, and deaminase (Figure 4b, Table 1). Most importantly, these are the first EATs demonstrating antibacterial activity. Seven EATs include DUF4157 domains as part of the protein or in a nearby gene (Table 1).

**Figure 4.**
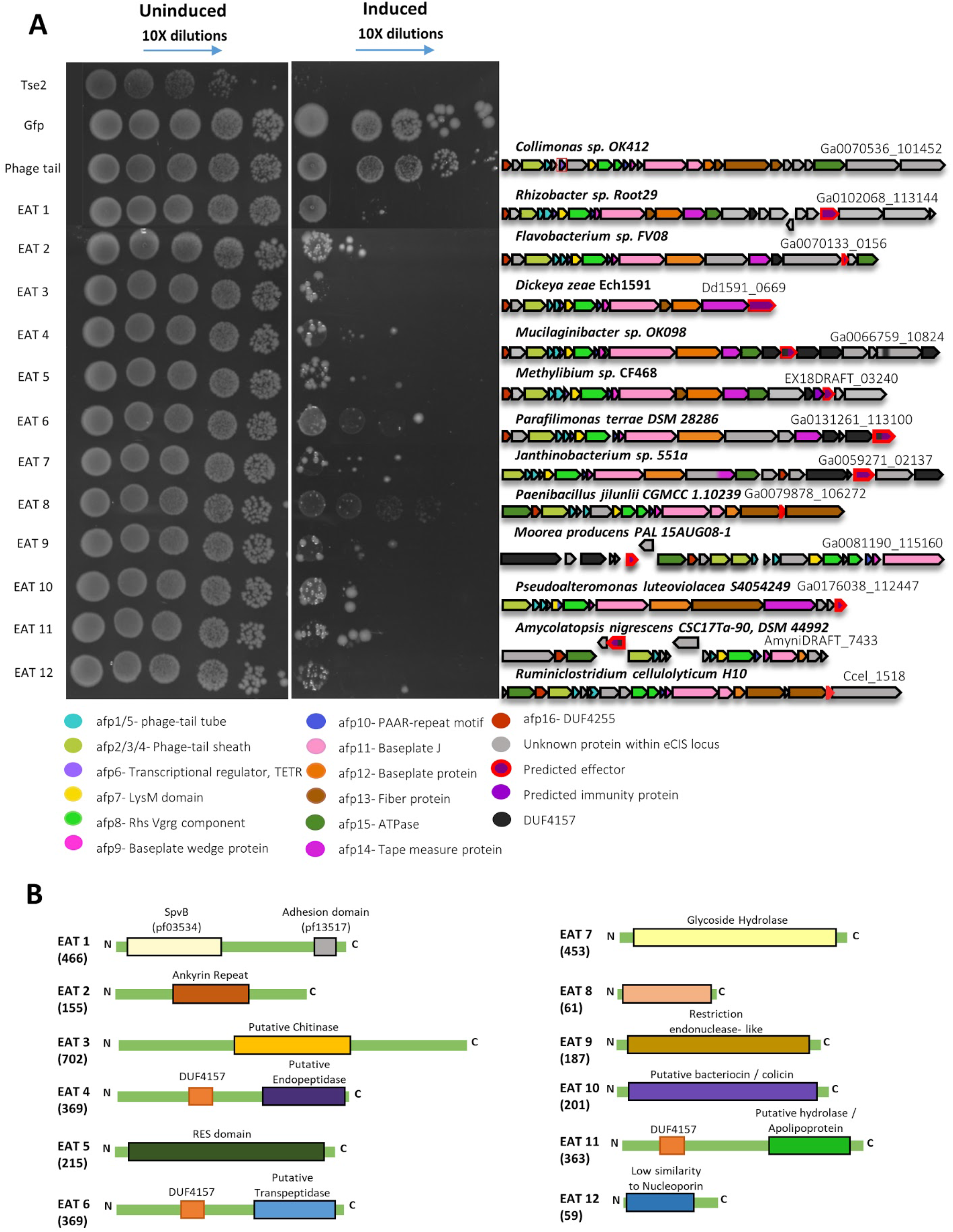
EATs are toxic to *E. coli.* EATs were cloned into pET28/29, transformed into *E. coli,* and were serially diluted and plated onto agar in induction conditions (right panel) or non-inducing conditions (left panel). tse2, a known Type VI Secretion System effector, was used as a positive control. Two negative controls used were meGFP, and an eCIS gene (phage tail). On the right side of each experiment is a cartoon representing the operon of the eCIS which the EAT was cloned from (purple gene with a red border); below is a legend explaining each component’s color. Image shows representative results of one experiment out of 2 experiments with similar results. **B.** Schematic representation of EAT protein and the domains within them. Size in amino acids is indicated in parentheses.

**Table 1.** New eCIS-associated toxins (EATs) experimentally validated in the current study.

| Toxin Name | Gene ID (IMG gene id) | Organism (taxon) | Activity | Predicted Molecular Function | DUF4147 gene/protein association |
| --- | --- | --- | --- | --- | --- |
| EAT1 | Ga0102068_1<br>13144<br>(2644141250) | *Rhizobacter* sp.<br>Root29<br>(Betaproteobacteria) | Antibact.,<br>anti-euk. | ADP-ribosyl<br>transferase<br>(SpvB) and<br>adhesion<br>(VCBS) | Absent |
| EAT2 | Ga0070133_0<br>156<br>(2616189889) | *Flavobacterium* sp.<br>FV08 (Bacteroidetes) | Antibact. | Unknown.<br>Encodes<br>ankyrin<br>repeats. | One and two<br>genes<br>upstream to<br>EAT2 |
| EAT3 | Dd1591_0669<br>(644851057) | *Dickeya zeae*<br>Ech1591<br>(Gammaproteo.) | Antibact.,<br>anti-euk. | Chitinase? | Absent |
| EAT4 | Ga0066759_1<br>0824<br>(2609591113) | *Mucilaginibacter* sp.<br>OK098<br>(Bacteroidetes) | Antibact. | peptidyl-Lys<br>metalloendo<br>peptidase | Within protein<br>center |
| EAT5 | EX18DRAFT_0<br>3240<br>(2587734256) | *Methylibium* sp.<br>CF468<br>(Betaproteobacteria) | Antibact. | NAD+<br>Phosphorylas<br>e | Two genes<br>upstream to<br>EAT5. |
| EAT6 | Ga0131261_1<br>13100<br>(2695001213) | *Parafilimonas terrae*<br>DSM 28286<br>(Bacteroidetes) | Antibact. | L,D-transpep<br>tidase | Within protein |
| EAT7 | Ga0059271_0<br>2137<br>(2602024578) | *Janthinobacterium*<br>sp. 551a<br>(Betaproteobacteria) | Antibact. | Unknown | Within protein<br>(N terminus) |
| EAT8 | Ga0079878_1<br>06272<br>(2668039404) | *Paenibacillus jilunlii*<br>CGMCC 1.10239<br>(Firmicutes) | Antibact. | Unknown | Absent |
| EAT9 | Ga0081190_1<br>15160<br>(2631153083) | *Moorea producens*<br>PAL 15AUG08-1<br>(Cyanobacteria) | Antibact. | Restriction<br>endonucleas<br>e | Three genes<br>upstream to<br>EAT9. |
| EAT10 | Ga0176038_1<br>12447<br>(2720476818) | *Pseudoalteromonas*<br>*luteoviolacea*<br>S4054249<br>(Gammaproteo.) | Antibact. | Peptidoglyca<br>n hydrolysis | Absent |
| EAT11 | AmyniDRAFT_<br>7433<br>(2515141031) | *Amycolatopsis*<br>*nigrescens*<br>CSC17Ta-90, DSM<br>44992<br>(Actinobacteria) | Antibact.,<br>anti-euk. | Deaminase | Within protein<br>(N terminus) |
| EAT12 | Ccel_1518<br>(643608442) | *Ruminiclostridium*<br>*cellulolyticum* H10<br>(Firmicutes) | Antibact. | Nucleoporin? | Absent |
| EAT13 | Ga0111076_1<br>23119<br>(2653242303) | *Candidatus*<br>*Udaeobacter*<br>*copiosus*<br>(Verrucomicrobia) | Anti-euk. | Actin<br>cross-linking | Flanked by<br>genes with<br>DUF4157<br>domain |

Antibact. - toxins killing the bacterium *E. coli*. Anti-euk. - toxins killing the yeast *S. cerevisiae*.

### Several eCIS-associated toxins are genetically linked to antitoxins

We identified several cases of toxin-immunity gene pairs next to eCIS operons supporting the antibacterial role of these eCIS loci as they require immunity genes (antitoxins) to protect from self-intoxication (Figure 5A). In one case, The toxin, EAT5, resembles RES (Figure 4b), an NADase that is accompanied by an antitoxin called Xre ^50,51^. We identified a gene, EX18DRAFT_03239, encoding an unknown protein located upstream of the toxin. We predicted that it serves as a new cognate immunity gene for the RES family, as the gene pair organization is conserved in phylogenetically diverse bacterial genomes (Figure S5). Expression of the RES-like EAT5 toxin was indeed toxic to *E. coli* while the putative immunity gene was not (Figure 5B). Co-expression of the toxin-immunity pair was sufficient to rescue bacteria from toxicity (Figure 5B). Finally, we showed that an intact immunity protein is required for the rescue phenotype, since deletion of 17 amino acids which encompass the C terminal alpha helix domain from the antitoxin abolished the immunity function (Figure 5B). Interestingly, the tail fibre protein (encoded by gene EX18DRAFT_03233) of the associated eCIS resembles a protein from Thermus phage YS40_Isch (see gene 2587734249 at Table S6), providing additional evidence that this eCIS targets bacteria and is therefore accompanied by an immunity protein.

**Figure 5.**
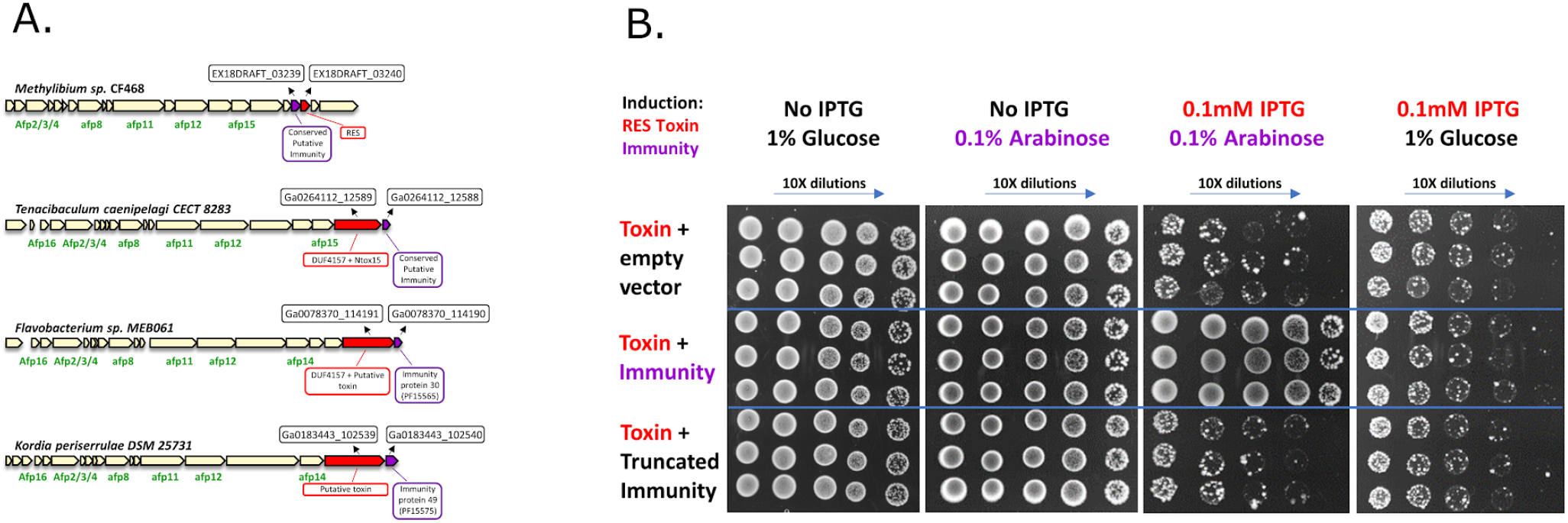
Toxin-immunity pairs associated with eCIS. **A. Pairs of known antibacterial toxins found in eCIS**. eCIS operons containing known or putative antibacterial toxic or immunity domains. **B. XRE-like Antitoxin Protects From RES-like Toxin.** Induction by IPTG leads to expression of RES-like toxin (IMG locus tag EX18DRAFT_03240). Glucose represses XRE-like antitoxin (IMG locus tag EX18DRAFT_03239) or its truncated version while Arabinose induces expression of XRE-like antitoxin (or its truncated version). Each section (separated by blue lines) represents three biological repeats.

Puffing the data together, we present here the first 12 antibacterial EATs and the first EAT-immunity gene pairs.

### Experimental validation of novel four eCIS toxins that target eukaryotic cells

Following our extensive bioinformatic analysis we were also interested in discovery of novel eCIS anti-eukaryotic toxins. We predicted new EATs that target eukaryotes based on their presence in the 3’ end of the eCIS operon, presence of eukaryotic domains within the encoded proteins, predicted enzymatic activity on eukaryote-specific molecules (such as actin), and similarity to known virulence factors. We cloned 14 genes into pESC-leu galactose-inducible vectors and transformed them into *S. cerevisiae* cells, serving as models for eukaryotic cells (Materials and Methods). Upon expression induction we identified four EATs that efficiently killed yeast (Figures 6, S9). Three EATs, EAT1, EAT3, and EAT11, also killed *E. coli.* EAT13 gene was tested only in yeast. These are predicted to act as an ADP-ribosyl transferase, a chitinase, a deaminase, and an actin cross-linker (Table 1). We think that yeast is probably not the native target for many other anti-eukaryotic EATs and therefore, it is likely that some of the toxins we predicted may be active when expressed in different organisms found in the native habitat of eCIS-encoding microbes.

**Figure 6.**
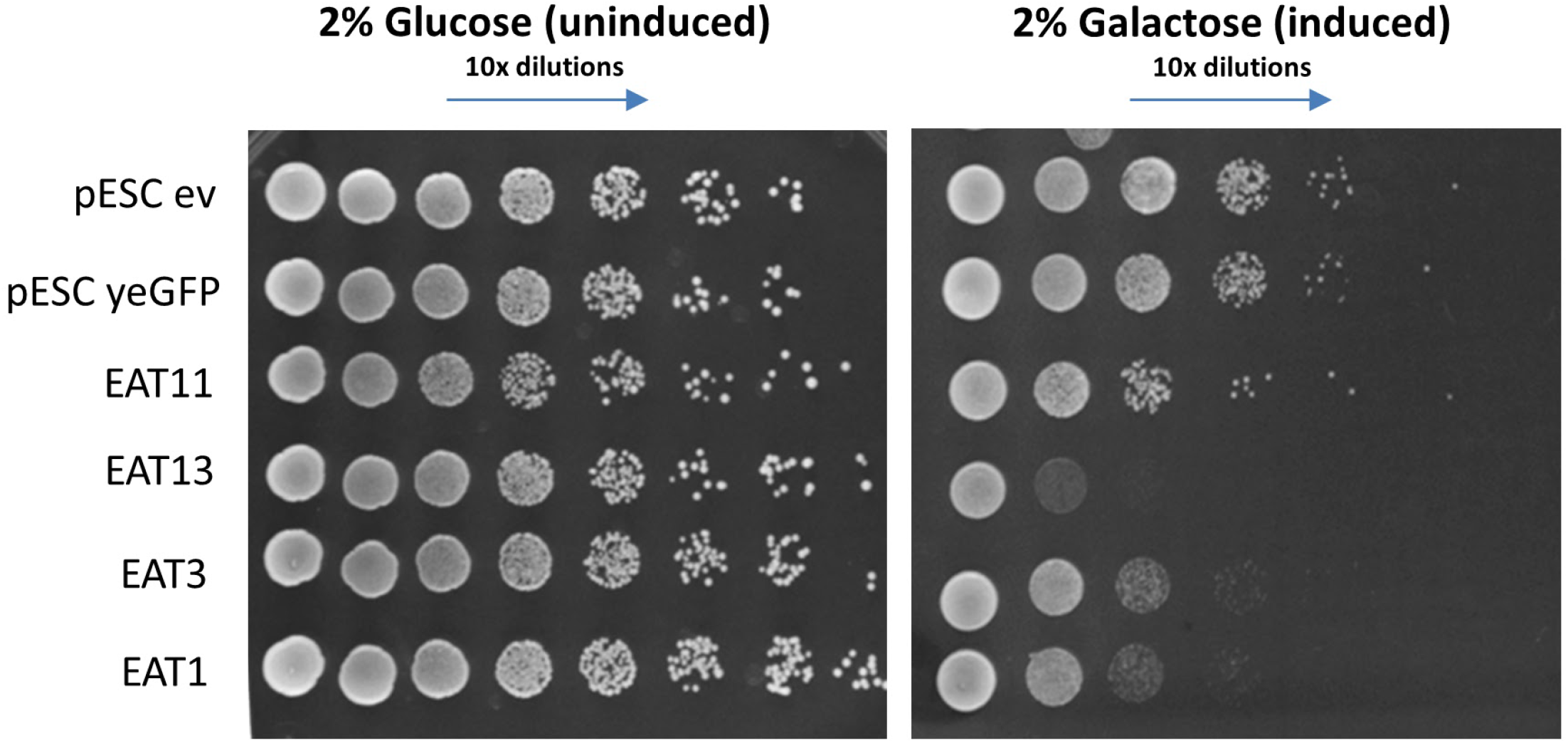
EATs are toxic to *S. cerevisiae.* EATs were cloned into pESC -leu Galactose inducible plasmids that were then transformed into Saccharomyces Cerevisiae BY4742 strain. Overnight cultures of the strains harboring the vectors of interest were grown in SD -leu media. The cultures OD was normalized and then washed once with water and split into two: one part was grown overnight in repressive conditions (SD -leu + 2% glucose) and the other part was grown in inductive conditions (SD -leu + 2% galactose). Dilutions were spotted on SD -leu plates containing glucose or galactose and the plates were incubated two nights at 30C. Negative controls: empty vector (ev) and non-toxin (yeGFP gene). Three biological replicates of this experiment are presented in Figure S10.

### Development of a comprehensive eCIS database

Finally, we developed a new repository called eCIStem that contains 1,425 operons from 1249 bacterial and archaeal genomes. The repository is publicly available at: http://ecistem.pythonanywhere.com. eCIStem provides visualization of all eCIS operons and includes gene annotation, an eCIS operon search option based on taxonomy, pfam domains, and gene clusters, and links to additional gene and protein data from the IMG database ^52^. eCIStem also contains ecological and life-style metadata of the eCIS encoding microbe. This valuable information can help researchers to study eCIS with regard to its specific ecological functions and to deduce the eCIS target organism. Figure 7 presents some of the information held in eCIStem. eCIStem is a significant expansion of the recently developed dbeCIS which includes 631 operons ^16^. We also expanded dbeCIS functionality by providing protein domain data for all eCIS genes, encoding organism metadata, and a full database download option which will facilitate future bioinformatic analysis of our dataset by the entire

**Figure 7.**
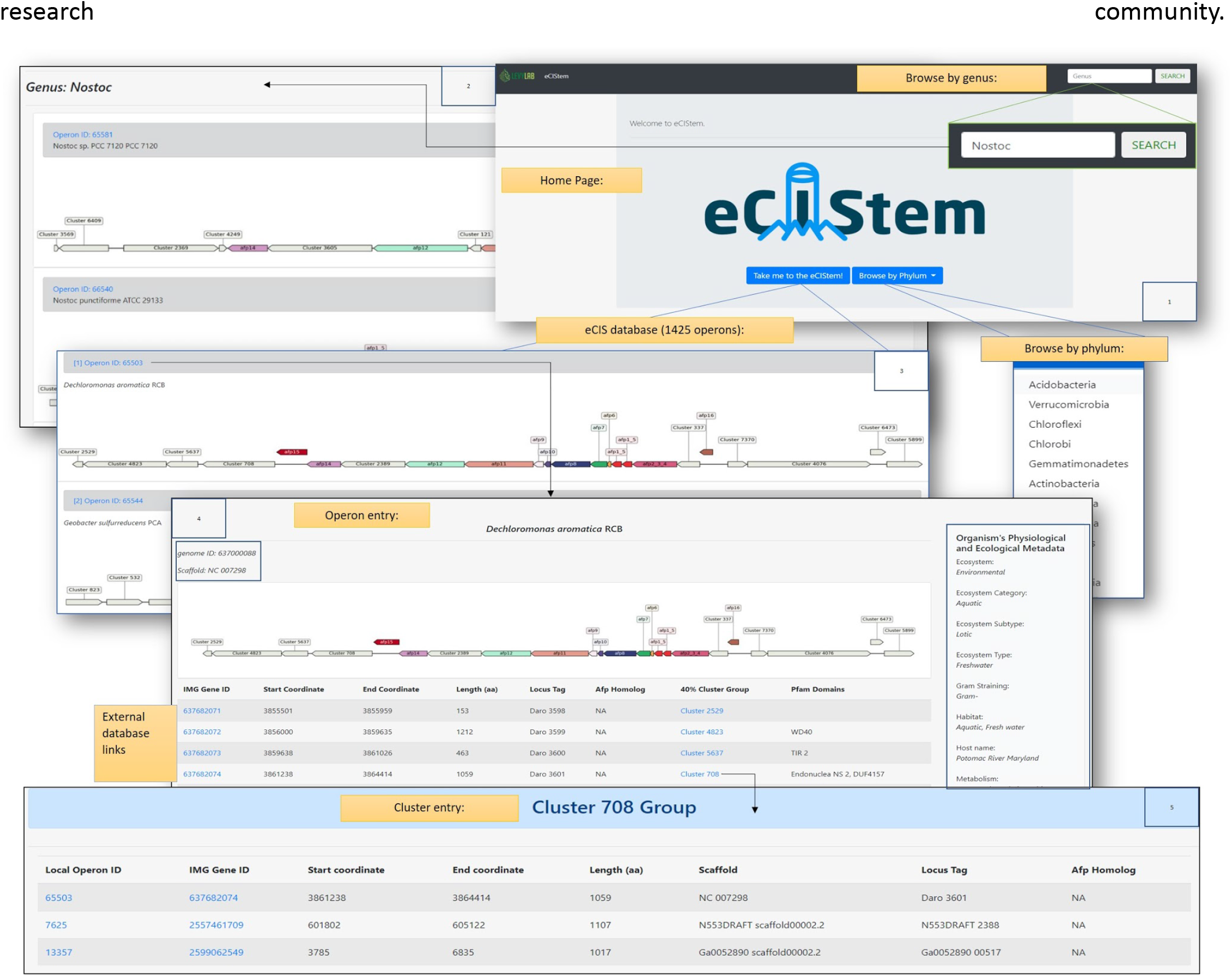
eCIStem database provides information on 1425 eCIS operons and their encoding genomes. Blue arrows represent links from page to page. **A.** A view of eCIS gene operons. **B.** An operon entry with information of each gene in the operon with operon, gene, protein, and protein domain information. **C**. Examples of eCIS protein cluster (clustered by 40% amino acids sequence identity). **D**. Examples of pfam domains entry for a specific gene. The information held in eCIStem is freely available to users from academic non-profit institutions.

## Discussion

In this study we use bioinformatic, phylogenetic, and molecular methods in order to thoroughly characterize the eCIS system across the microbial world. Based on our results we provide an improved model for eCIS function that enriches our understanding of eCIS ecological and taxonomic distribution within microbes, potential eCIS regulation and cell attachment mechanisms. Moreover, we expanded the known repertoire of eCIS targets. We show that, most likely, eCISs not only target eukaryotic cells, as previously reported, but also bacteria (Figure 8). Our extensive analysis uncovered new conserved eCIS core and shell components that encode the eCIS structure and regulation and numerous EATs genetically associated with eCIS loci. The high number and variety of EATs suggest that eCIS loci can serve as a treasure trove for toxin gene discovery. Previous works uncovered seven EATs, all from Gammaproteobacteria class, whereas here we added 13 novel EATs from six different bacterial phyla. We examine the occurrence of eCIS operons within a large-scale genome set and provide a strong support for eCIS enrichment in environmental microbes from soil and aquatic ecosystems and hosts, and a depletion from mammalian and avian microbiomes; with a specific depletion from their pathogens.

**Figure 8.**
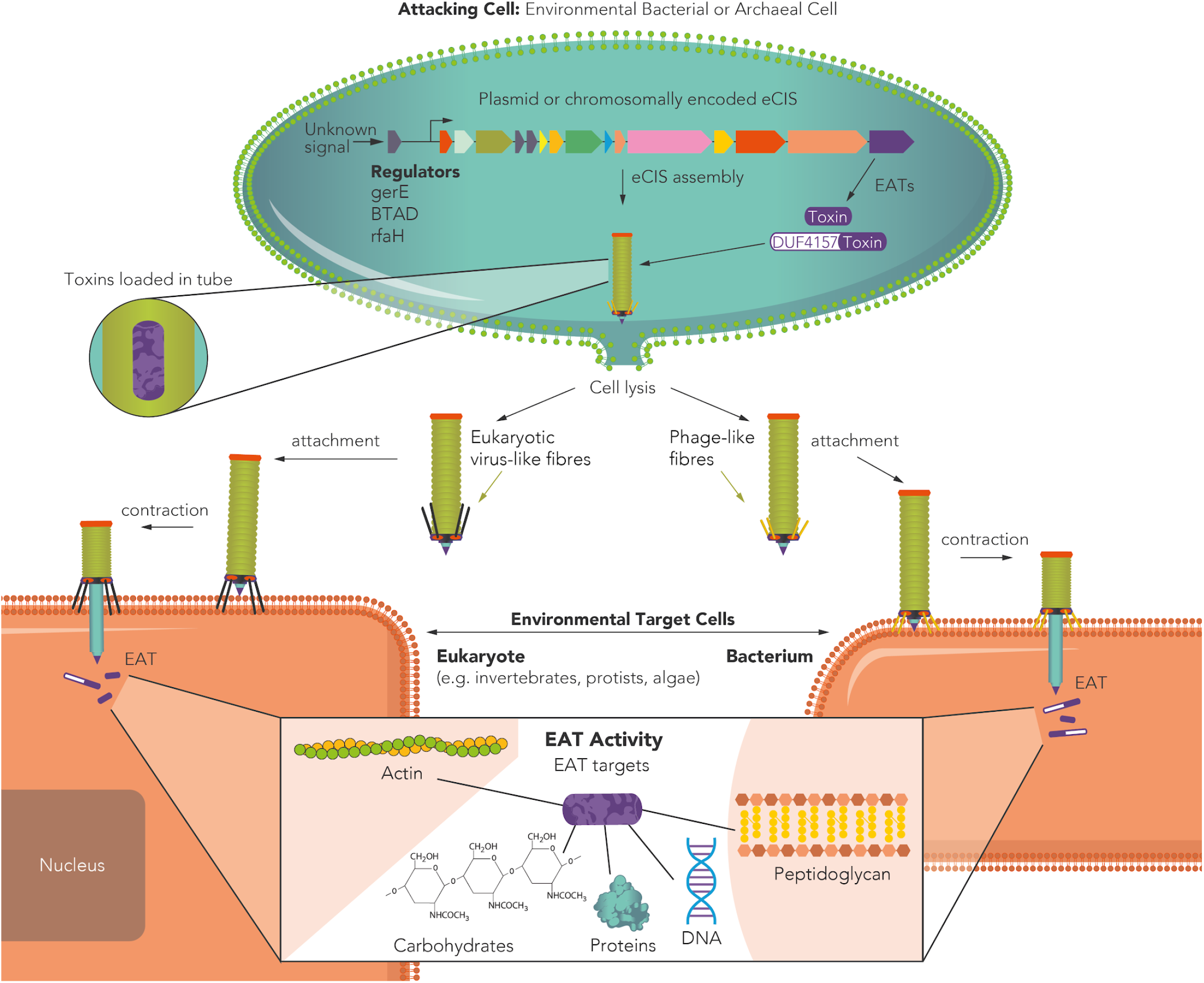
A revised model for eCIS function and ecology. eCIS is encoded by operons that are enriched in environmental microbes. These operons are regulated by adjacent genes such as RfaH ^77^ and genes that carry GerE and BTAD domains. The regulators are occasionally activated by kinases and are activated/repressed by yet unknown signals. Effectors (EATs) tend to be encoded in the 3’ end of the operons. EATs often carry DUF4157 domains in their N-terminus or found adjacent to genes with this domain. The eCIS particle is assembled with the toxins inside the tube ^14^ and is released following cell lysis ^12^. The particles carry tail fibres that mediate attachment to environmental eukaryotic or prokaryotic target cells. The tail fibres attach, contract, and release the EATs inside the target cells, mostly leading to its death or growth arrest. The EATs function intracellularly in diverse ways, enzymatically acting on different cellular components that were identified in this work and by other groups.

We identified that some eCIS tail fibres that can help predict whether eCIS target bacteria or eukaryotes. Interestingly we identified eCIS tail fibres that share high sequence similarity with similar proteins of *Yasminevirus* giant virus and *Phycodnavirus,* which infect amoebae ^53^ and algae ^54^, respectively (Figure S4). Based on this finding we propose that eCIS from these bacteria could target similar hosts. The fact that most eCIS tail fibers matched both phage and eukaryotic-targeting viruses suggests they may have broad spectrum binding activity, and therefore broad targeting activity.

One intriguing mystery is related to the antibacterial EATs and their lack of immunity genes in most cases. Nearly all antibacterial toxins produced by bacteria are accompanied by adjacent immunity genes ^36,39,55–61^, which we could not bioinformatically identify next to most of the newly verified EAT genes. We do not know how these antibacterial EATs are being produced and loaded onto the eCIS system without killing the producer cell prior to eCIS release. One possibility is that these are produced right before the cell is lysed to release the eCIS, and therefore the cell does not require protection at this time, as it is already “fated” to die, and cellular activity shifts to releasing eCIS particles. It is possible that EATs even purposely contribute to lysis. Another option is related to the yet mysterious function of the DUF4157 domain which accompanies many of the EATs. This domain may confer some immunity to the eCIS producers through temporal toxin inactivation.

Shedding light on eCIS biological role in nature can reveal new means by which microbes affect the health and development of their hosts and shape neighboring eukaryotic and prokaryotic microbial communities.

## Materials and Methods

### Identification of eCIS operons within microbial genomes

The starting database for identification of eCIS was 64,756 publicly available genomes from the Integrated Microbial Genomes database ^52^, which includes all protein-coding genes along with their pfam annotations, as per the IMG pipeline ^62^. Genomes were searched for the presence of genes annotated as containing pfam domains ^63^ associated with eCIS (Pfam IDs: 00004, 04865, 04965, 04984, 05488, 05954, 06841, 14065). Genes encoding proteins with the aforementioned pfam annotations were grouped by physical linkage (encoded within 12,000 base pairs of one another). Only those groups of linked genes with at least four members, with at least three of them having different pfam annotations, were considered putative eCIS operons. Because of the evolutionary similarity between eCIS and other contractile machinery, we removed any putative operons with genes annotated with phage-related pfams (Pfam IDs: 04860, 03237, 17289, 05929, 05125, 05944, 05926, 04550, 05840, 04606, 05065, 07068, 16855, 06810, 03864, 02924, 10124), such as capsid genes, within 10,000 base pairs of any genes in our putative operons. Furthermore, we removed any putative operons with genes annotated as COGs ^64^ associated with Type VI Secretion Systems (COG IDs: 0542, 3521, 3523) within 10,000 base pairs of any genes in our putative operons. We believe this is a very stringent filtration protocol that was also added to filtration of pyocins (see below). The boundaries of these putative operons were expanded by 10 genes up- and downstream of the extrema of the putative operons to identify eCIS accessory (non-core) genes. Only protein coding sequences were kept, and tRNA and other genes were ignored. Using previously published HMM profiles of eCIS core elements ^16^, and the hmmsearch tool (a part of the HMMER package; ^65^), the expanded putative operons were surveyed and their genes were annotated as core elements. Only the top half of the hmmsearch results were used (according to bitscore), in order to minimize false positive hits. Furthermore, genes annotated by IMG as having pfams that match core elements were also annotated as core genes. The core-annotated putative operons were filtered for those with at least 10 core genes, and with at least one eCIS-specific core (AFP 12, 13, 14, or 16; ^16^), in order to remove any possible R-type pyocins from the dataset. The final database of eCIS operons contains 1,249 genomes, with 1,425 distinct operons.

### Phylogenetic eCIS gene tree construction

Amino acid sequences for Afp11 and Afp8 homologs were retrieved from each eCIS operon. In the case of operons with more than one copy of either the Afp8 or Afp11 homologs, the genes with the larger sequence were chosen. In the minority of cases (15/1425) where one or both of the genes were missing, they were not included in the tree. The Afp11 and Afp8 sequences were separately aligned with clustal omega ^66^ using standard seffings. The alignments were trimmed using TrimAl ^67^ with the “automated1” flag selected. The Afp11 and Afp8 alignments from corresponding operons were concatenated. The concatenated alignments were inputted into FastTree using standard seffings ^68^, and visualized and annotated using interactive Tree Of Life (iTOL; ^69^).

### Systematic tail fibre (Afp13) sequence analysis

629 Afp13 amino acid sequences, as defined by genes labeled by hmmsearch as afp13 within the top 25th percentile of bitscores, were inputted as queries for two searches using BLASTP against the NCBI nr database using standard parameters ^70^. In the first search, the nr database was filtered by taxon IDs corresponding to tailed bacteriophages (NCBI taxid: 28883). In the second search, the nr database was filtered by taxon IDs corresponding to eukaryotic-targeting viruses, by manually removing prokaryotic viruses from taxid 10239 (see Table S6 for full taxid list). The results from each search were filtered for those hits with p-value < 0.001, leaving 296 unique Afp13 hits. Each query Afp13 gene was checked for the hit with the top bitscore from the first and second searches to determine if a given gene better aligned with tailed phages, or with eukaryotic-targeting viruses (Table S6). Those Afp13 with bitscore difference greater than 15 were marked as having a preference for one dataset over another; within 15 was considered inconclusive.

### Detection of eCIS within plasmids

Together with our collaborator Dr. William Anreopolous (San Jose State University) we developed Deeplasmid (https://sourceforge.net/p/deeplasmid/). We ran Deeplasmid on all scaffolds carrying eCIS. Deeplasmid uses a deep learning algorithm that classifies scaffolds as being chromosomes or plasmids based on different features, such as GC content, characteristic genes, and coding density. The algorithm achieves an AUC-ROC of over 93% based on our tests. The score cutoff of >= 0.7 was chosen as it correctly identifies the pADAP plasmid that encodes the Antifeeding Prophage (AFP). Other known plasmids carrying eCIS received Deeplasmid scores higher than 0.95 (Table S3).

### eCIS association with genera

The taxonomic makeup of the eCIS database was compared to the number of genera in publicly available genomes of Bacteria, Archaea, and plasmids of bacteria from IMG using a custom python script. A Fisher exact test was calculated for each genus, which returned an odds ratio and an associated p-value. Multiple hypothesis testing correction was performed using the Benjamini Hochberg procedure. Those genera with a q-value of less than 0.001 and at least 10 entries in the publicly available IMG database are displayed in Figure 1.

### eCIS association with ecological and physiological features

To correct for taxonomic bias that affects enrichment analysis, a population-aware enrichment analysis based on Scoary ^27^ was used. Scoary accepts input of a tree to analyze Fisher exact test enrichments in the context of phylogeny. To create the tree, 16s sequences for each genome with eCIS was queried. In the case of more than one 16s gene per genome, one was chosen randomly. These sequences were inputted into clustal omega ^66^ to generate a UPGMA guide tree. The inputs to Scoary were (1) the guide tree, (2) a presence-absence file, indicating if a genome has or lacks an eCIS operon, (3) metadata files that indicate for each genome if they possess a certain characteristic, e.g. whether it was isolated from soil. This program outputs an odds ratio, a q-value for the Fisher exact test that produced the odds ratio, and a q-value based on how many contrasting pairs (instances not from the same clade) support the enrichment, i.e. how many examples of the enrichment are independent of taxonomy.

### Protein Domain (Pfam) Enrichment Analysis

We collected the pfam annotations for the 64,756 proteins encoded by publicly available genomes from IMG. Based on our eCIS operon database, we marked all the pfam domains that were found in our eCIS database as “pfam domains within eCIS operons”. Duplicated domains within a single gene (usually of repeat domains) were dropped, in order to avoid inflation of enrichment. We then counted all the occurrences for each pfam - within eCIS operons, outside eCIS operons, and in total. After counting all pfam occurrences we performed a Fisher Exact Test for pfam enrichment in eCIS operons, compared to the rest of genomes in the analysis. Multiple hypothesis testing correction was performed using the Benjamini Hochberg procedure. The adjusted P value (q value) of the Core Component was zero, so in order to plot it we changed it to 1e^−250^. Then we plotted the data using R ^71^. We counted the number of Phyla each Pfam domain appeared in eCIS. Pfam domains appearing in >10 Phyla were marked as “Core Domains”, domains found in 4-10 Phyla were marked as “Shell Domains”, and domains found in <4 were marked as “Cloud Domains”. These terms were borrowed from pangenomics.

### Toxins and accessory gene clustering

To identify accessory genes, i.e. genes that are not annotated as homologs of Afp proteins, the operons were expanded from their extrema by four protein coding genes up- and downstream. All genes lacking an annotation of an Afp homolog were clustered. This included the genes up and downstream of extrema, as well as genes in between annotated Afp homologs. Using CD-HIT ^72^, the amino acid sequences of these unmarked genes were clustered with a threshold of at least 40% identity over at least 80% of the length of both query and subject. Each gene was inputted into CD-Search ^73^ with standard seffings to determine if the gene encoded for conserved domains. Each 40%-identity-protein-cluster was then manually examined for domains associated with known biological function, e.g. known toxin domains, known transcription factor domains, etc.

### Heterologous expression of putative EATs in bacteria

Candidate gene sequences were retrieved from IMG ^52^ and were synthesized, following codon adaptation, and cloned into either pET28, pET29, or pBAD24 plasmids by Twist Bioscience. The plasmids were then transformed into *E. coli* BL21 (DE3) pLysS strain. Overnight cultures of the strains harboring the vectors of interest were grown in LB containing the proper selection: Kan or Amp for pET28/29 or pBAD24, respectively. The cultures were normalized by OD to be equivalent and subsequently serially diluted. Dilutions were spotted on LB agar containing the proper selection and inducer (500 μM IPTG, 0.1% arabinose) or repressor (1% glucose) and the plates were incubated overnight at 37C.

The toxin-antitoxin assay was done as followed: E. coli BL21 (DE3) pLysS strain was transformed with the toxin (EAT 5) on a pET28 plasmid and its predicted antitoxin (IMG gene id 2587734255) on a pBAD24. Two control strains were made, an empty vector (pBAD24) and a truncated antitoxin. We removed the last 51 nucleotides (17 amino acids) from the gene (which was synthesized by TWIST bioscience) using inverse PCR, then it was re-ligated with Nebuilder (NEB) kit. All three strains were tested in a drop assay with three biological replicates. Induction of toxin and antitoxin was performed with 100 μM IPTG and 0.1% arabinose respectively.

### Heterologous expression of putative toxins in yeast

Candidate sequences were retrieved from IMG ^52^. They were synthesized, following codon adaptation to yeast by Twist Bioscience and were cloned into pESC -leu Galactose inducible plasmids. The plasmids were then transformed into *Saccharomyces Cerevisiae* BY4742 strain ^74^. Overnight cultures of the strains harboring the vectors of interest were grown in SD -leu media. The cultures OD was normalized and then washed once with water and split into two: one part was grown overnight in SD-leu 2% glucose media and the other part was grown in SD-leu 2% galactose media. Dilutions were spotted on SD-leu plates containing glucose or galactose and the plates were incubated for 48 hours at 30C.

### Construction of eCIStem repository

The ecis database was constructed based on the eCIS data from our research (Tables S1, S2, S7, S8). The information is divided into four tables: Operons Table, Gene Table, Organism Table and Pfam Table. The eCIStem database scheme is described in Figure S11. The Django framework was used to represent the data and its relationships and deploy it for public usage.

### S74 Peptidase (PF13884) secondary structure analysis

We clustered all 84 genes containing PF13884, using CD-HIT with 40% similarity ^72^. Used two cluster representatives for structural prediction in Phyre2 ^75^, and secondary structure alignment using Praline ^76^.

## Acknowledgments

AL is generously supported by the Israeli Science Foundation (Grant #1535/20) and Alon Fellowship of the Israeli council of higher education. AMG is generously supported by the Kaete Klausner Scholarship and a scholarship from the Israeli Ministry of Aliyah and Integration. We thank Prof. Maya Schuldiner, Prof. Yechiel Shay, and Prof. Saul Burdman for providing different plasmids and cells. We thank Dr. Omri Finkel, Dr. Hila Sberro, Prof. Saul Burdman, Prof. Rotem Sorek, Dr. Erez Mills, Dr. Gal Ofir, and Dr. Lianet Noda-Garcia for careful evaluation of the manuscript.

## Supplementary Figures

**Figure S1.**
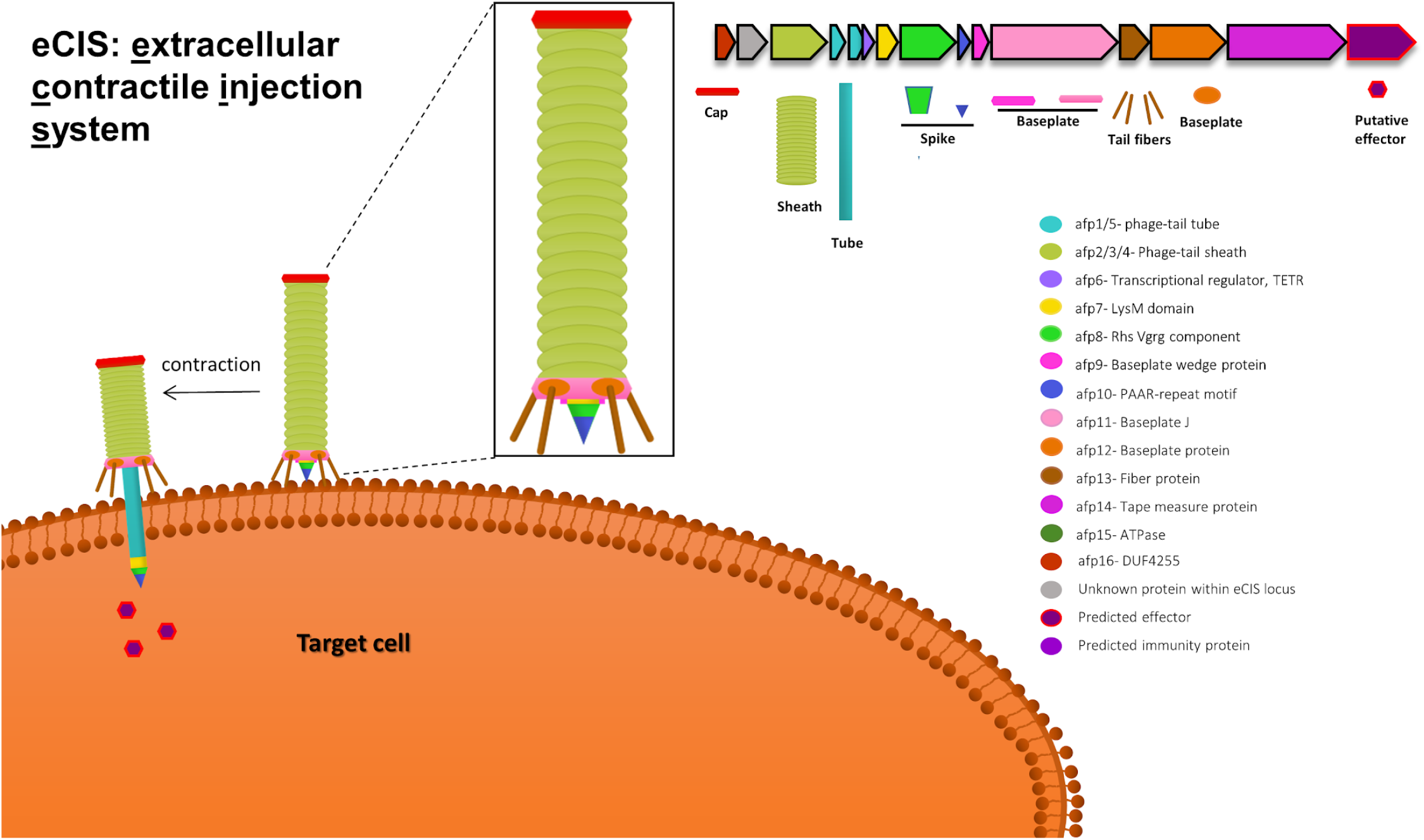
eCIS structure. eCIS particles are encoded by an operon of 15-28 genes, a typical operon is shown, with cartoons showing corresponding structural components (right side). The core proteins encode a sheath that has a cap on one side and a baseplate on the other side. Inside the sheath there is a tube that ends with a sharp spike. The particle likely binds to receptors located on the target cell through tail fibres. Once the contact is established, the eCIS contracts, injects the tube, and eCIS-associated toxins (EATs) are released into the target cell (left side). Some of the toxins were reported to be located inside the hollow tube. The illustration is based on Jiang et al. ^3^

**Figure S2.**
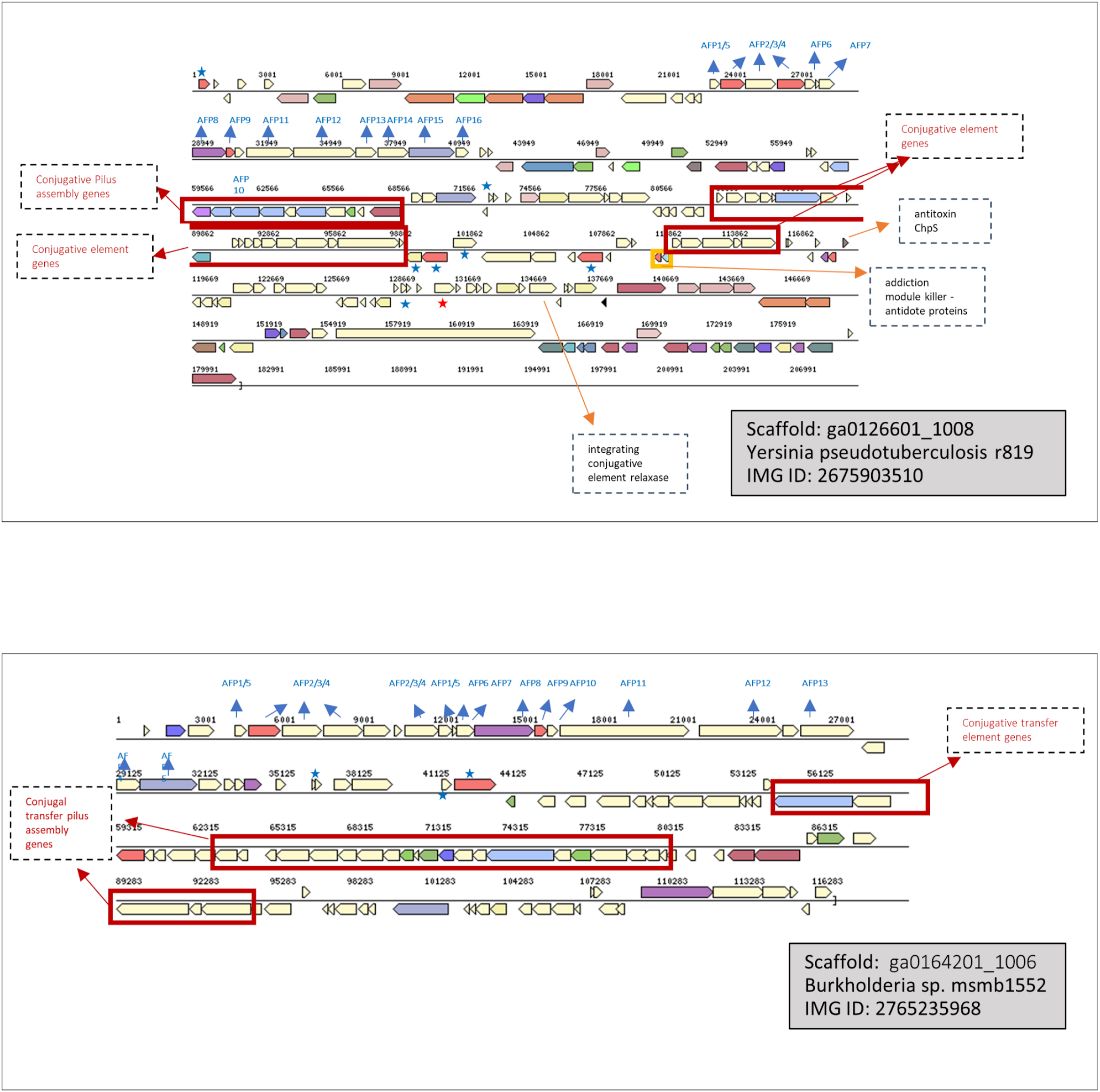
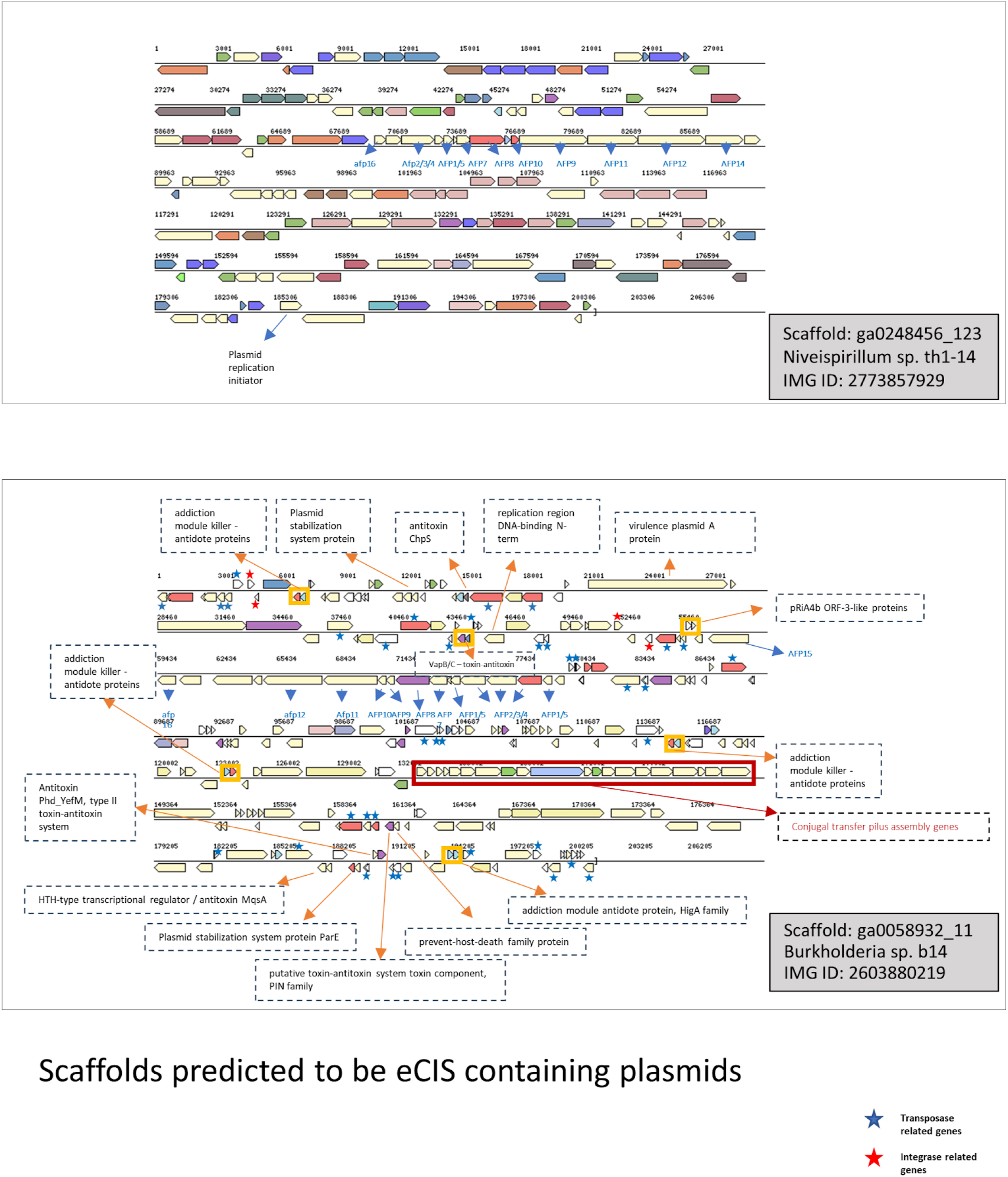
Four examples of predicted plasmid-borne eCIS operons. Genes that play a role in plasmidic activities are marked.

**Figure S3.**
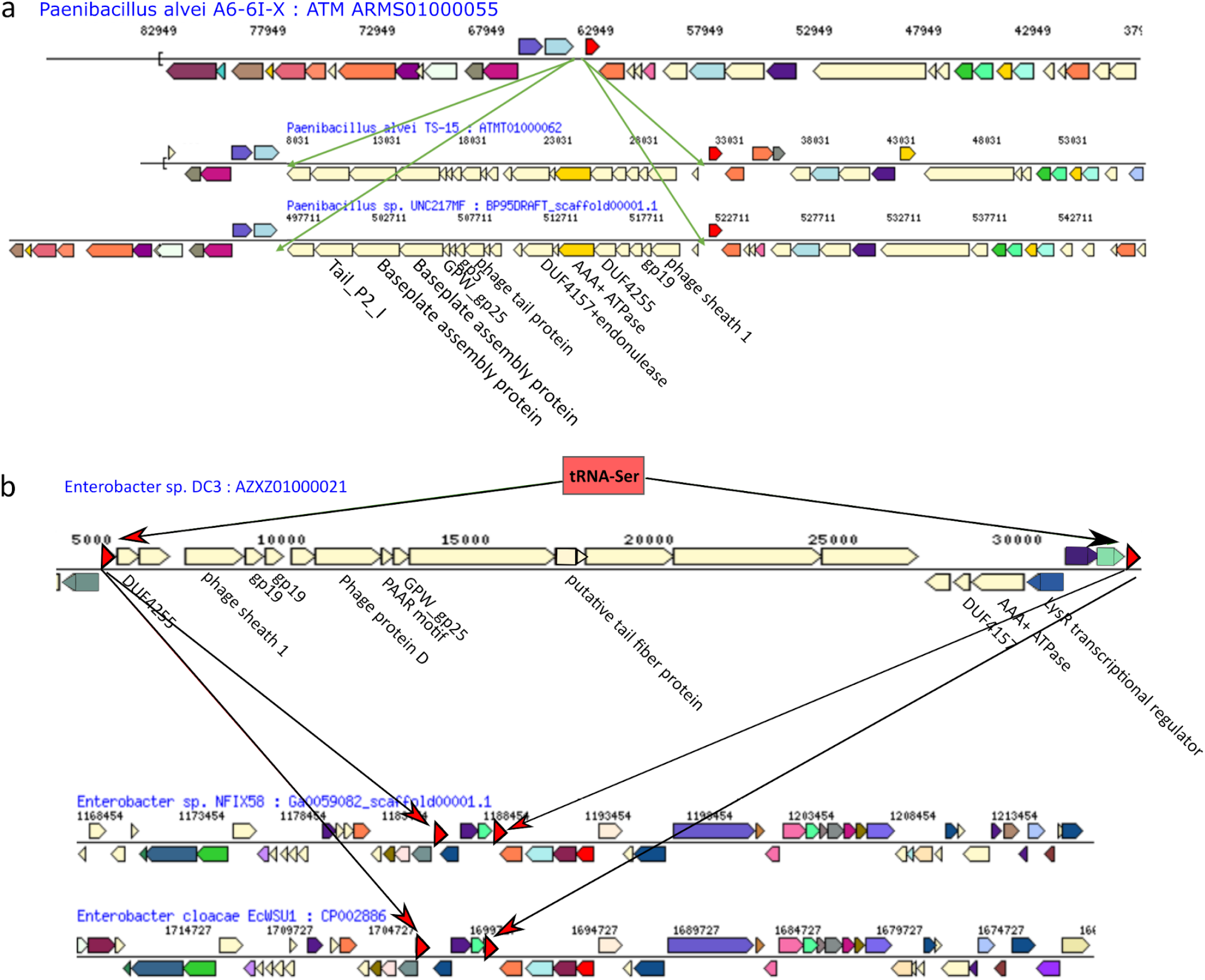
Evidence for local eCIS locus integration events into bacterial genomes. **a.** A local eCIS integration into specific strains of *Paenibacillus* strain. **b.** A local eCIS insertion into an *Enterobacter* strain. The integration was likely mediated via homologous recombination using the flanking eCIS tRNA-Ser genes and an external eCIS copy that carries this gene.

**Figure S4.**
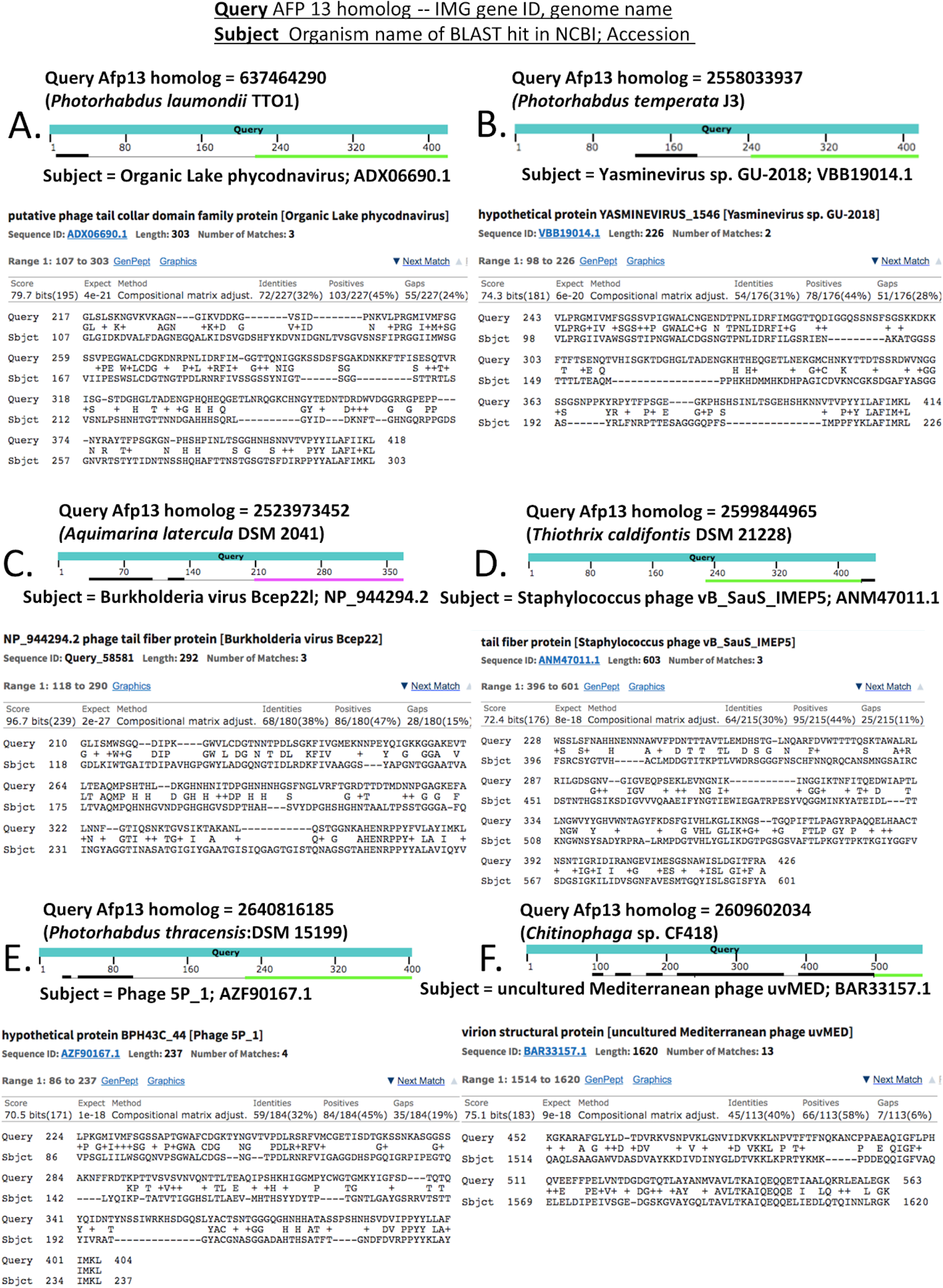
Examples of high identity BLAST protein sequence alignments between Afp13 sequence queries and tail fibre proteins from bacteriophages and viruses as the targets.

**Figure S5.**
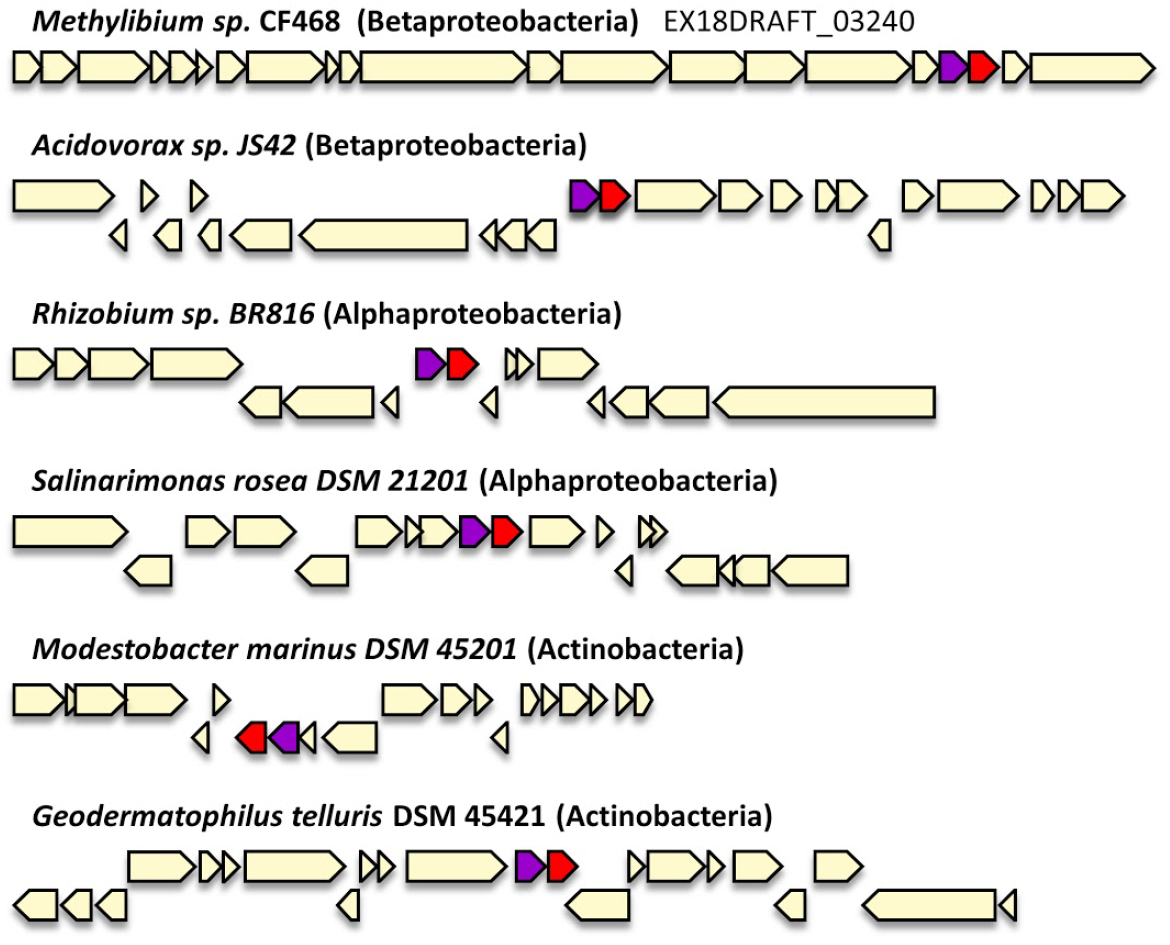
RES-Xre Gene pair syntenny. A gene pair containing the gene for EAT5 is conserved through various strains from proteobacteria and actinobacteria. All occurrences except for the top one are outside eCIS genomic context (IMG locus tag: Ajs_1583; RhiBR816DRAFT_0322;Swit_5317;Ga0056099_04803 ;Ga0056090_4110). Based on this syntenny we predicted that the purple gene serves as an antitoxin.

**Figure S6.**
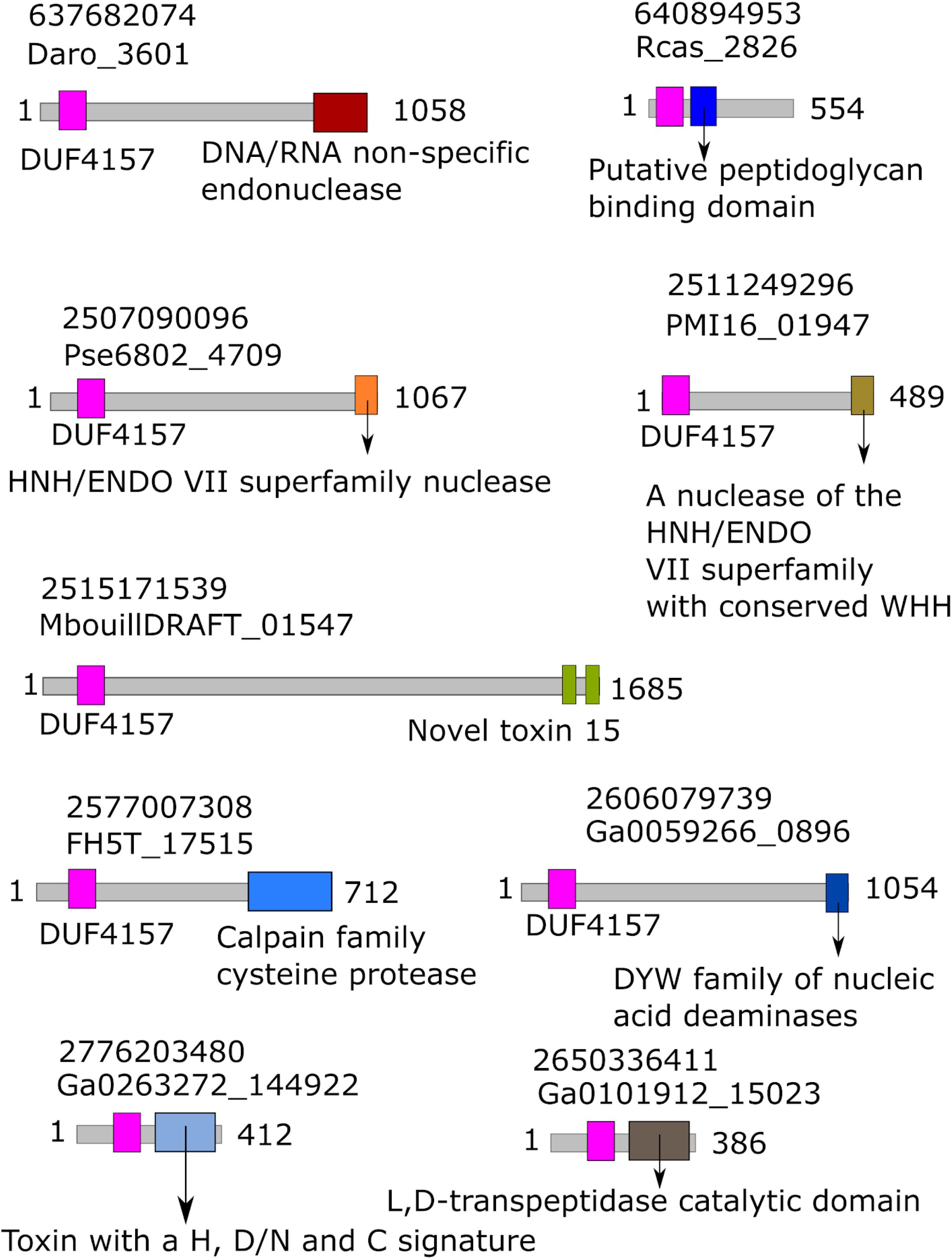
Examples for putative eCIS-associated toxins (EATs) carrying DUF4157 at the N terminus. The upper number indicates IMG gene ID. The lower number indicates the locus tag (from IMG website).

**Figure S7.**
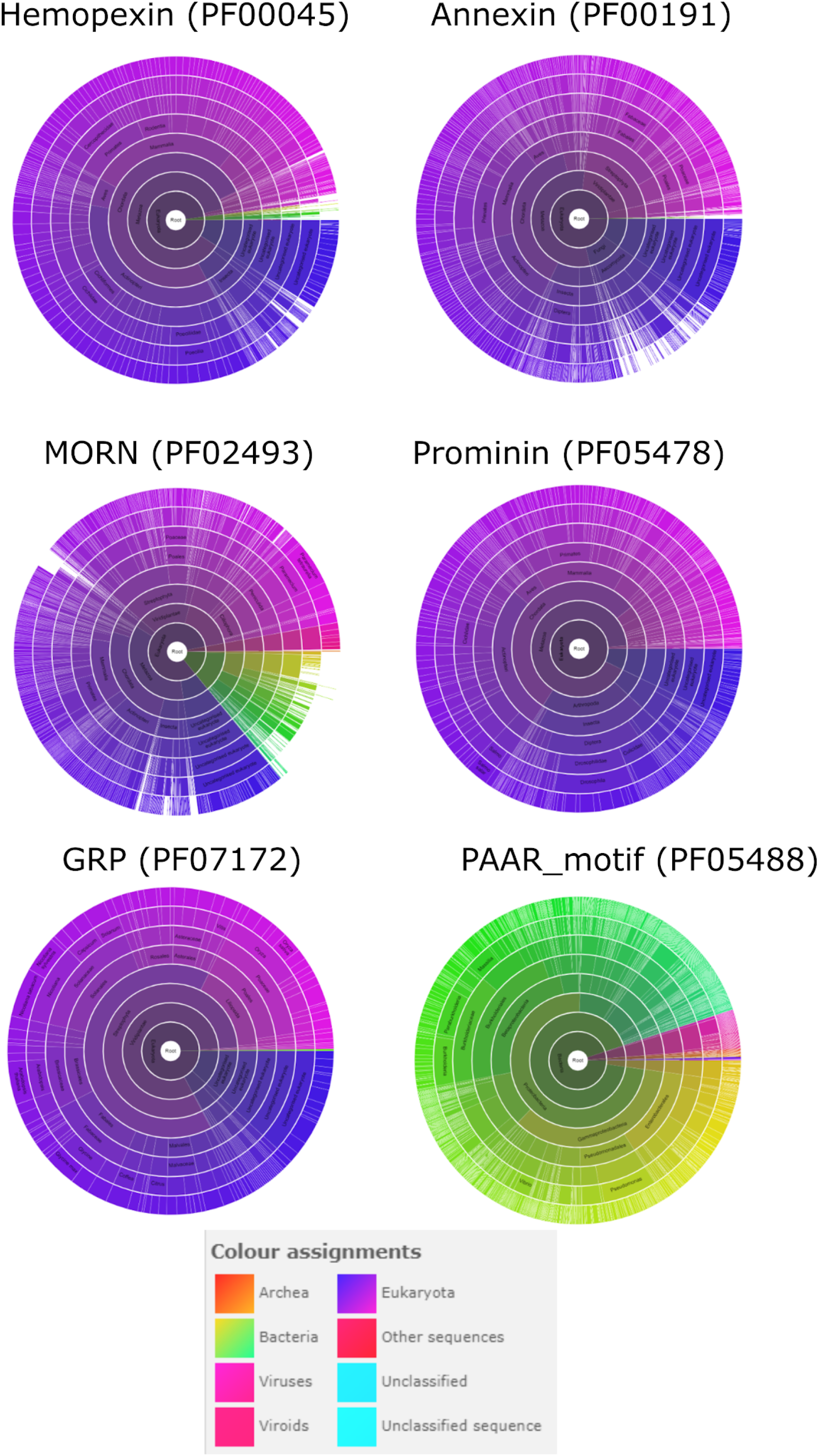
Several eCIS toxins carry protein domains that are mostly found in eukaryotic annotated genomes. The sunburst plots, representing species distribution and taken from Pfam website https://pfam.xfam.org/, represent the distribution of protein domains in available annotated genomes. The color coding for each plot appears below. As a reference we added the PAAR motif found in the eCIS spike which is mostly a bacterial domain.

**Figure S8.**
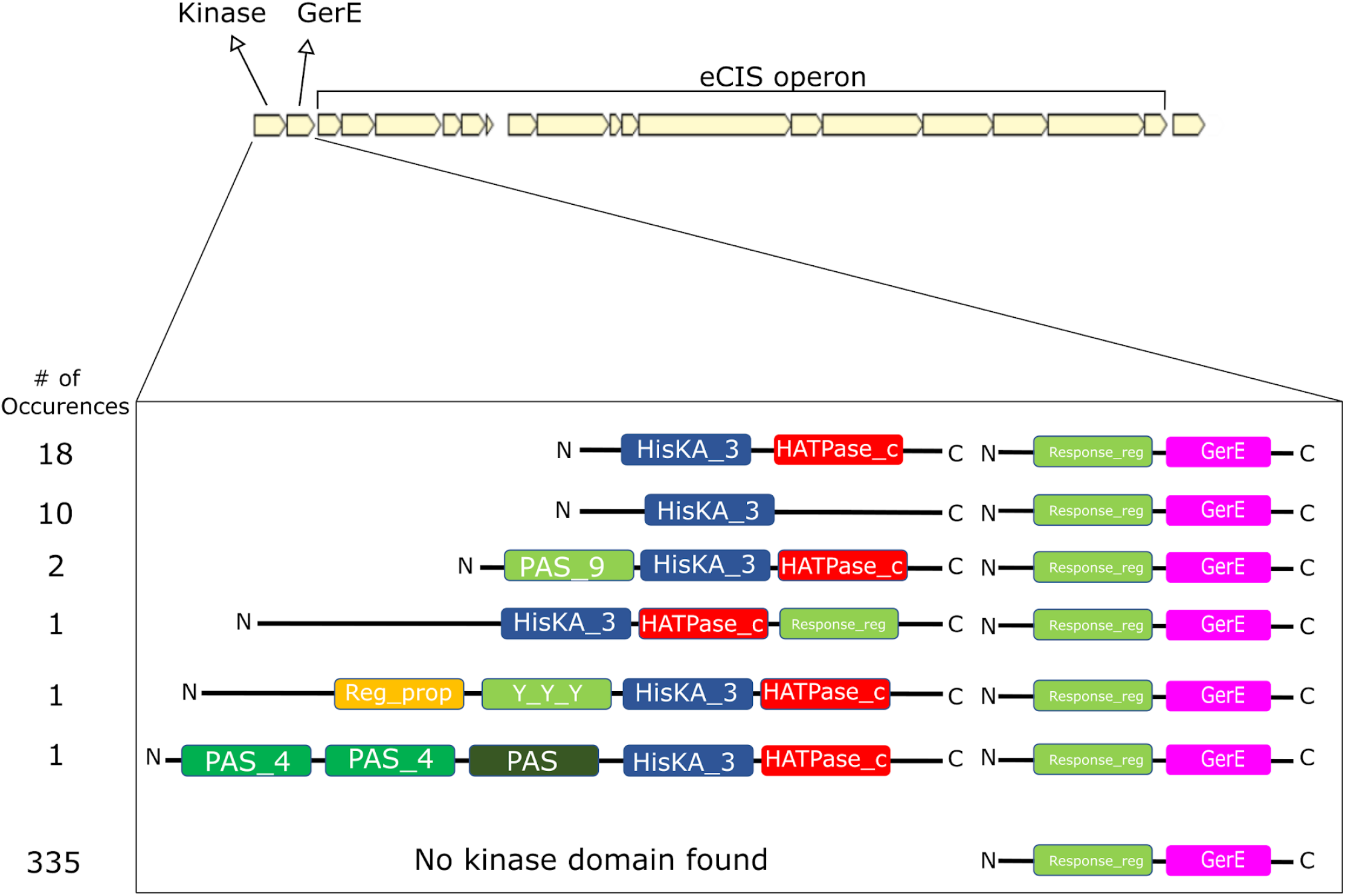
Kinase domains are found upstream to the putative GerE eCIS transcription regulator. An illustration of the different pfam domain architectures and their occurrences within the kinase genes found upstream to the GerE gene that is found in the eCIS operons.

**Figure S9.**
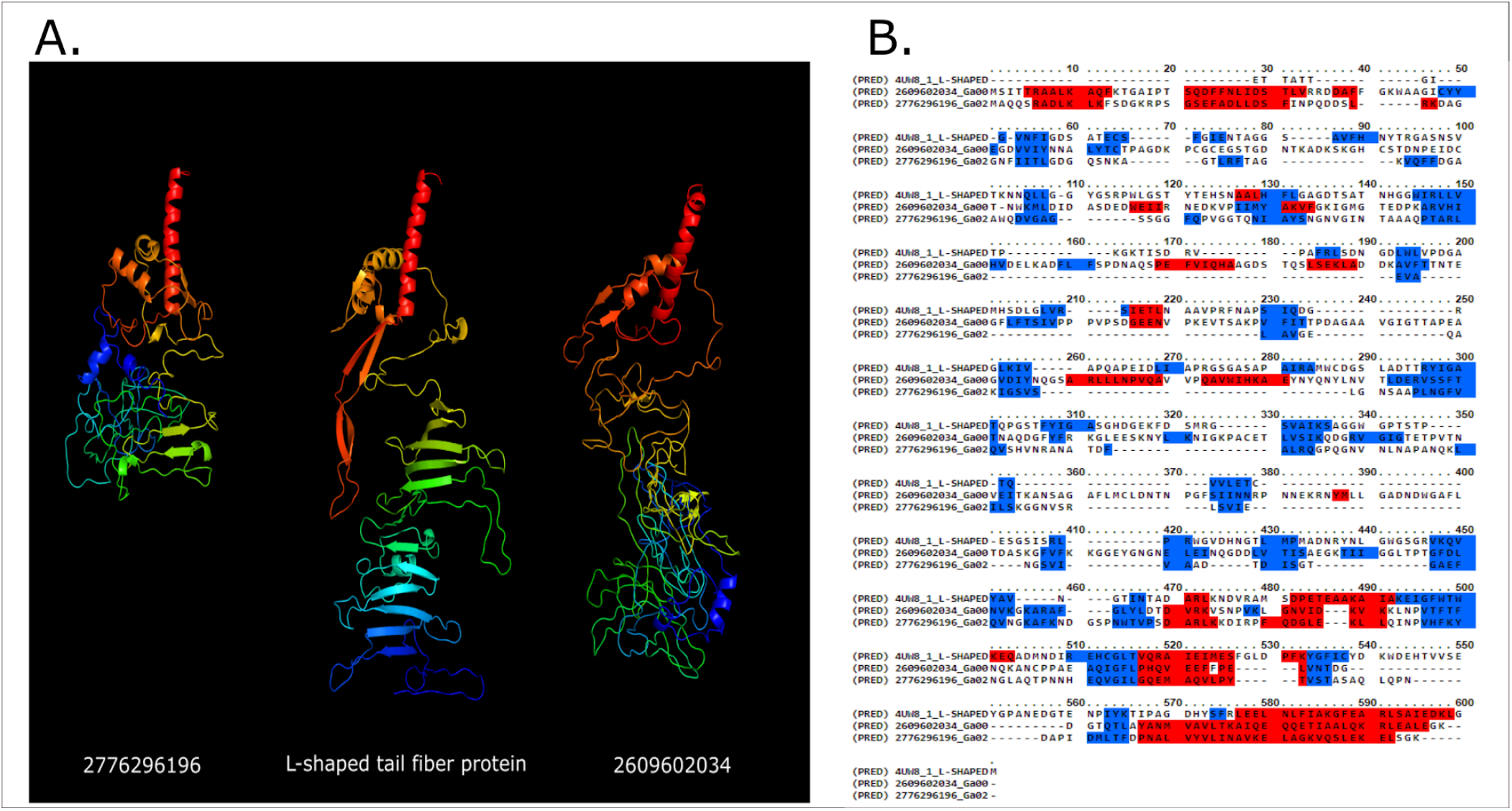
Genes containing S_74 peptidase domain structurally resemble the carboxy-terminal domain of the T5 Phage tail fiber. **A.** 3D structure of cluster representatives (IMG: 2609602034, 2776296196) containing the S_74 Peptidase domain, compared to the carboxy terminal domain of the T5 tail fiber (PDB accession: 4UW8), predicted by Phyre2 with intensive modelling mode^75^. **B.** Alignment of the secondary structure with the T5 phage L-shaped tail fiber (PDB accession: 4UW8) using Praline ^78^.

**Figure S10.**
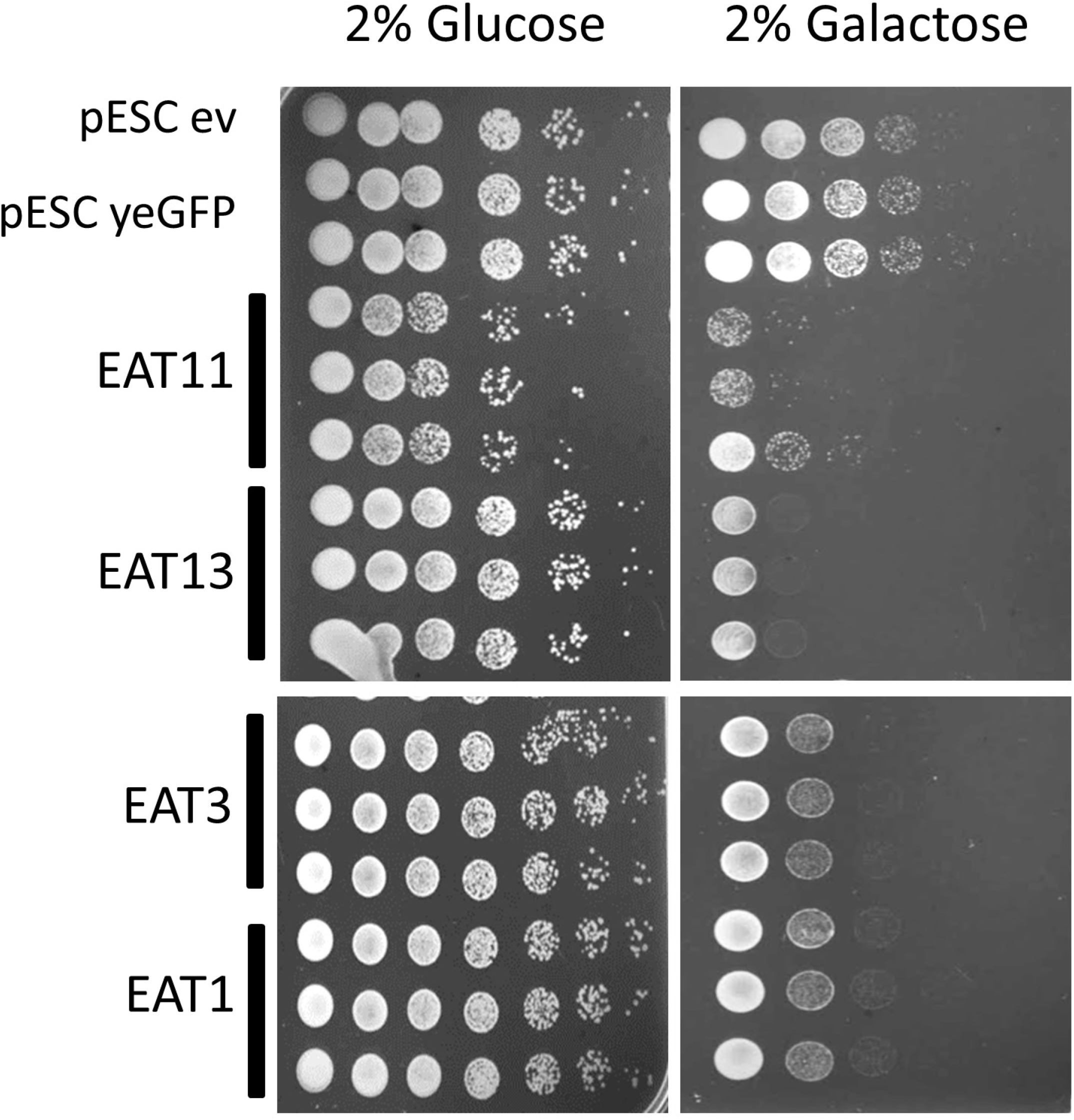
Biological replicates of EATs killing *S. cereviciae.* EATs were cloned into pESC -leu Galactose inducible plasmids that were then transformed into *Saccharomyces cerevisiae* BY4742 strain. Overnight cultures of the strains harboring the vectors of interest were grown in SD -leu media. The cultures OD was normalized and then washed once with water and split into two: one part was grown overnight in repressive conditions (SD -leu + 2% glucose) and the other part was grown in inductive conditions (SD -leu + 2% galactose). Dilutions were spotted on SD -leu plates containing glucose or galactose and the plates were incubated two nights at 30C. Negative controls: empty vector (ev) and non-toxin (yeGFP gene).

**Figure S11.**
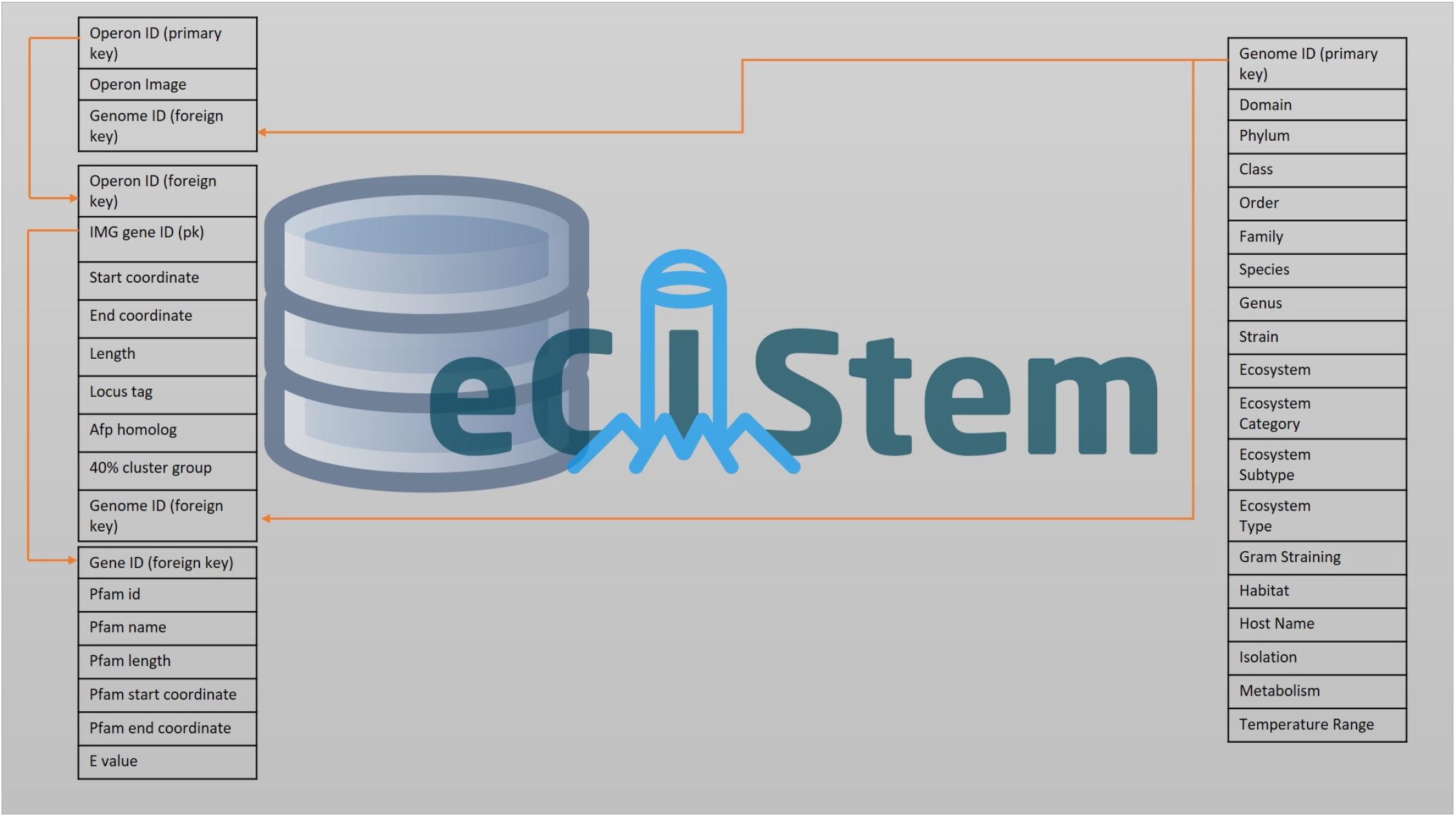
eCIStem database scheme.

## Supplementary Tables

**Table S1. eCIS operons.** Information about the 1426 eCIS operons we identified and their encoding organisms.

**Table S2. Genes included in eCIS operons**.

**Table S3. eCIS predicted to be carried on plasmids.** We ran Deeplasmid tool on all contigs/scaffolds carrying eCIS operon and annotated plasmids based on Deeplasmid score >0.7. Deeplasmid is a deep learning based classifier that identifies plasmids based on presence of plasmidic genes and other features such as coding density, and GC content.

**Table S4. Correlation between eCIS and genera.** For each genus in our genomic database we performed a fisher exact test by counting the number of sequenced genomes within the genus encoding eCIS, those that lack eCIS, genomes from all other genera that encode eCIS, and enomes from all other genera that lack eCIS.

**Table S5. eCIS correlation with microbial ecological and physiological features.** All data was downloaded from IMG database.

**Table S6. BLAST results of Apf13 proteins against proteins from viruses and phages.** The best hits are presented.

**Table S7. Pfam enrichment in eCIS operons.** Results of Fisher exact test for enrichment of the different Pfam domains within proteins encoded by eCIS operons.

**Table S8. eCIS Accessory Gene Clusters.** This table shows all accessory proteins (i.e. non-AFP homologs), clustered by at least 40% identity over at least 80% of each of their lengths. Cluster ID = a unique nominal identifier for each 40% identity group. Blank entries in Conserved Domain columns represent no hit from the NCBI CDD database. Widespreadness refers to each cluster, and how many of the indicated taxa are represented in each cluster. Operon ID eCIStem Database is a unique identifier for the eCIStem database (see full text for more information).

**Table S9. Core Gene Clusters.** This table shows clusters of core genes (i.e. AFP homologs). Each cluster is a unique nominal identifier for genes with at least 40% ID over at least 80% of clusters.

## Notes

### Competing Interest Statement

The authors have declared no competing interest.

